# Widespread low-affinity motifs enhance chromatin accessibility and regulatory potential in mESCs

**DOI:** 10.1101/2025.11.18.685822

**Authors:** Melanie Weilert, Kaelan J. Brennan, Khyati Dalal, Sabrina Krueger, Haining Jiang, Rosa Martinez-Corral, Julia Zeitlinger

## Abstract

Low-affinity transcription factor (TF) motifs are an important element of the cis-regulatory code, yet they are notoriously difficult to map and mechanistically incompletely understood, limiting our ability to interpret non-coding variation in development, evolution, and disease. Here we investigate their role in pioneering and leverage sequence-to-profile models of chromatin accessibility in mouse embryonic stem cells to reliably map and interpret low-affinity motifs across the genome. We find that low-affinity motifs have outsized effects by cooperating with nearby motifs through intra-nucleosomal soft syntax. By modeling nucleosome-mediated cooperativity with a kinetic model, we discover and validate that pioneer cooperativity makes a motif operate at higher pioneering ranges across changing TF concentrations, thereby raising the regulatory potential. These results show that low-affinity motifs can be accurately mapped, shape the properties of developmental enhancers and likely play a widespread role in fine-tuning enhancers during evolution.

## Introduction

One of the unresolved fundamental problems in biology is the cis-regulatory code, which, if mapped and understood, could unlock the gene regulatory information encoded in the human genome.^1,2^ Mutations in transcription factor (TF) binding motifs can cause phenotypic changes, making it necessary to obtain comprehensive maps of all functional motifs across cell types to predict the effect of non-coding genetic variants.^3^ A key challenge is to map low-affinity motifs, which are often critical for the specificity of enhancers during development.^4–10^ These motifs allow an enhancer to selectively become active when a certain combination of TFs are present, and their mutation can cause phenotypic effects.^6,7^

Mapping functional low-affinity motifs across the genome is however challenging. Since they are defined by weak TF binding strength *in vitro* and low match scores to a position weight matrix (PWM), local sequence scanning produces too many false positives.^11–14^ Mapping low-affinity motifs therefore requires an understanding of the sequence context in which they become functional. Traditional mechanistic studies have mostly focused on DNA-mediated TF binding cooperativity, which occurs at close distances of less than ∼30 bp and can be examined with *in vitro* TF binding assays.^6,9,15–17^ How low-affinity motifs contribute to enhancer function beyond TF binding cooperativity has remained a gap in knowledge.

Here we studied the role of low-affinity motifs in pioneering—the first step in enhancer activation where the nucleosomal DNA is made accessible in chromatin.^18^ This process is carried out by pioneer TFs, which may cooperate^19,20^ or use low-affinity motifs.^19,21–23^ However, the effects vary widely across genomic regions and cell types, making it difficult to derive the underlying sequence rules.^24,25^ Furthermore, what constitutes a pioneer TF and what properties mediate its function are still debated, complicating efforts to examine the mechanisms of cooperativity. To make pioneering a tractable problem, we focus here on a single widely studied cell type, mouse embryonic stem cells (mESCs), and define a pioneer TF as any TF whose motif has a measurable effect on chromatin accessibility *in vivo*. This lens allows us to specifically assess the role of low-affinity motifs in pioneering.

An open question is whether low-affinity motifs confer specific properties to enhancers. Since low-affinity motifs are bound at higher TF concentrations,^10^ they allow developmental enhancers to respond at higher TF concentration ranges during embryonic development,^9,26–30^ making them susceptible to TF dosage-dependent phenotypic changes.^27^ However, alternative models have not been explored. For example, if low-affinity motifs rely on cooperativity with other motifs, this could also affect an enhancer’s properties. Yet, how to rigorously distinguish the effects of motif affinity from cooperative interactions across the genome to systematically identify the mechanisms by which enhancers function *in vivo* is not clear.

A major advance in mapping functional motifs in the genome and identifying their syntax rules has come from sequence-to-function deep learning models.^1,31,32^ These models are trained to accurately predict genomics data from DNA sequences and learn motifs combinatorially *de novo*, without prior biological assumptions. With post-hoc interpretation tools, the learned motif instances and syntax rules can be extracted.^33^ This approach has generated motif maps that explain TF binding,^34–36^ chromatin accessibility,^19,21,27,37–39^ reporter expression,^40,41^ promoter activity,^42–44^ and cell-type specific enhancer activity *in vivo*.^45–47^

While the predictive accuracy of the models is widely appreciated, the underlying molecular mechanisms that govern the functional outcomes of these syntax relationships are only beginning to emerge.^21,27,35,36,48–50^ Mechanistic interpretations of the cis-regulatory code learned by sequence-to-function models, including the role of low-affinity motifs, remain limited.

To investigate the role of low-affinity motifs in mediating pioneering in mESCs, we analyzed ChromBPNet models that predict bias-free chromatin accessibility at base-resolution from genomic sequences.^38^ These models accurately learn motifs that causally drive accessibility, and distinguish them from bystander motifs that are statistically enriched in the regions, but whose number, strength and location are not informative for the predictions.^21,38^ By interpreting these models, we quantify the contribution of individual TF motifs to pioneering, define the cooperativity syntax, and use the cooperative measurements to model the response to TF concentration changes.

We find that low-affinity motifs have substantial effects by cooperating with strong pioneer motifs within intra-nucleosomal distances (<200 bp). This syntax is far more flexible than the syntax of DNA-mediated TF binding cooperativity, suggesting a nucleosome-mediated mechanism. Modeling indicates that this cooperativity shifts a motif to higher pioneering ranges but in a way distinct from increasing the motif’s affinity. These results help explain why the sequence basis for pioneering has been difficult to decipher and suggest that low-affinity motifs play a widespread role in shaping the activity of developmental enhancers during evolution.

## Results

### Accessibility deep learning models enable the accurate mapping of low-affinity pioneer motifs in mESCs

To learn functional low-affinity motifs, we optimized existing deep learning approaches to train a ChromBPNet model that accurately predicts chromatin accessibility profiles in mESCs from DNA sequence^38,51^ (Figures 1A-B and S1A, Methods). We also trained a separate BPNet model on high-resolution TF binding data (ChIP-nexus) for Oct4, Sox2, Nanog, Klf4 and Zic3 in mESCs^36,51^ (Figures 1A and 1B) to independently discover, compare and validate the motifs of these TFs. We ensured that both models could learn the same motifs by training on the same 154,827 1 kb-long output genomic coordinates (Figure 1B, Methods), optimizing and cross-validating across different training set combinations (Table S1 and S2, Methods). Contribution scores were obtained using DeepSHAP, and sequences with high contribution towards accessibility or binding were clustered by TF-MoDISco into motifs, represented as either contribution weight matrices (CWMs) or or PWMs.^36,51–54^

**Figure 1.**
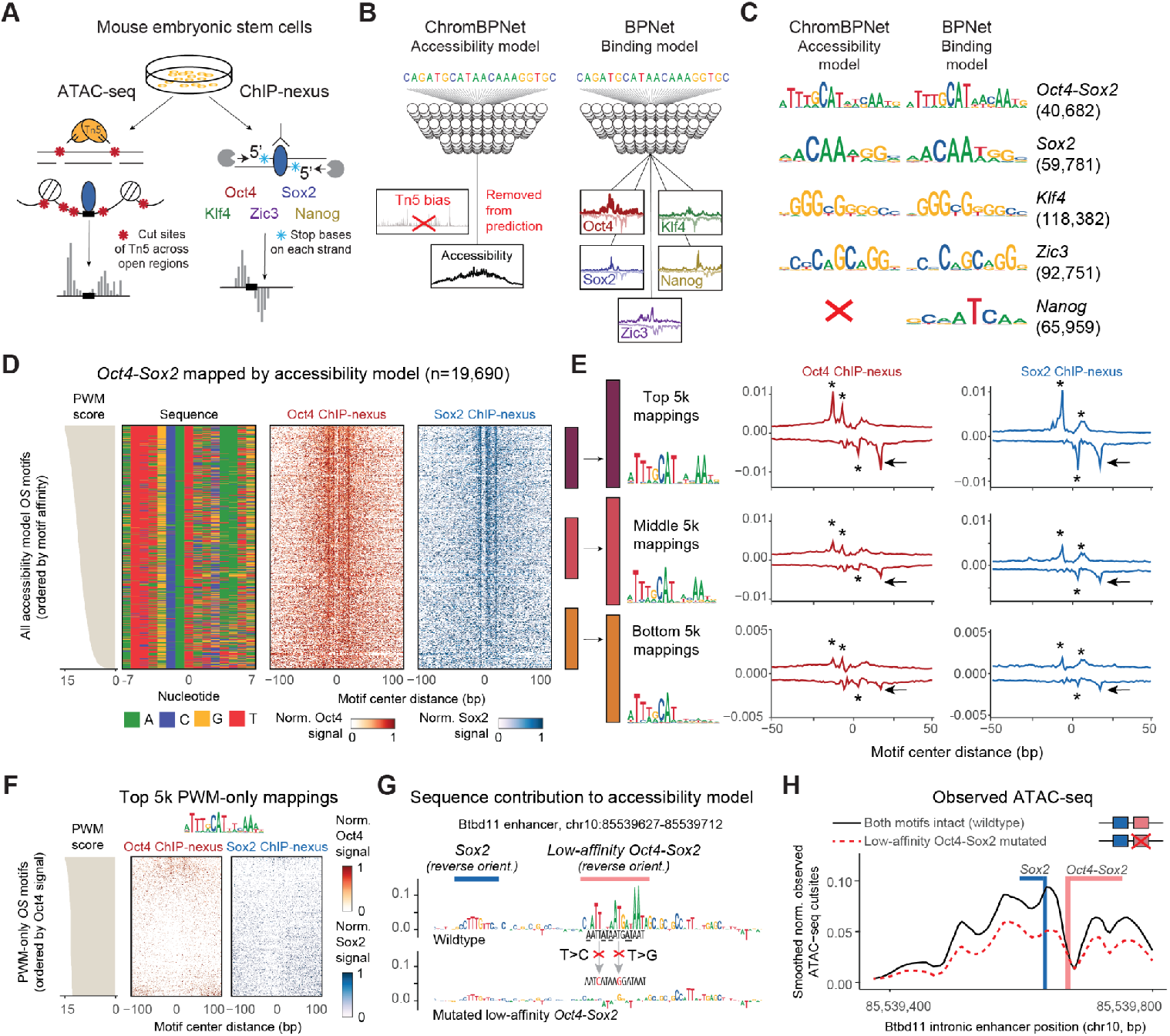
BPNet and ChromBPNet models predict and map functional low-affinity pioneer motifs. **A)** Graphic depiction of ATAC-seq and ChIP-nexus experiments performed on the mouse embryonic stem cell line R1. **B)** Graphic depiction of the sequence-to-profile models ChromBPNet and BPNet, trained to predict ATAC-seq and ChIP-nexus profiles of the pluripotency TFs Oct4, Sox2, Nanog, Klf4 and Zic3, respectively. **C)** The accessibility and binding models returned expected motifs for each of the five TFs, with the exception of *Nanog*, which was not found by the accessibility model. Motifs are represented as a contribution weight matrix (CWM) after combining motifs from both models. **D)** *Oct4-Sox2* motifs mapped by the accessibility model, ordered by position weight matrix (PWM) scores (left, beige) and shown as sequence colormap (second panel). The normalized heatmaps of non-stranded ChIP-nexus profiles for Oct4 (third panel, red) and Sox2 (fourth panel, blue), covering 200bp centered on the motifs, show that the majority of mapped motifs display both Oct4 and Sox2 ChIP-nexus footprints. **E)** The top, middle and bottom 5,000 *Oct4-Sox2* motifs based on the PWM scores are shown with their PWM logos (left) and alongside the average ChIP-nexus profiles of Oct4 (red) and Sox2 (blue). The y axis shows RPM-normalized reads on the positive strand (top) and negative strand (bottom); stars mark the ChIP-nexus footprints that demarcate the protein-DNA contacts on each strand. The black arrow marks the outer right ChIP-nexus footprint of Sox2 and Oct4, which persists even at the bottom 5k *Oct4-Sox2* motifs when the *Sox2* component is no longer clearly recognizable. **F)** Oct4 and Sox2 binding at the top 5,000 *Oct4-Sox2* motifs mapped <u>only</u> by PWM-scanning in the same regions, ordered by PWM scores (left, beige). Normalized heatmaps of the Oct4 (red) and Sox2 (blue) non-stranded ChIP-nexus profiles show very little signal recognizable as footprints despite the strong PWM match scores, showing that PWM scanning produces false positives. **G)** The intronic *Btbd11* enhancer shown with contribution scores from the ChromBPNet accessibility model across the wildtype sequence (top) and for the sequence where the low-affinity *Oct4-Sox2* motif was mutated by two base substitutions to abolish its accessibility contribution (bottom). Note that the low-affinity *Oct4-Sox2* motif (<u>A</u>ATT<u>A</u>T<u>A</u>ATG<u>AT</u>AAT) is the reverse complement of ATT<u>AT</u>CAT<u>T</u>A<u>T</u>AATT, which is the orientation of the motif introduced above and which has five mismatches (underlined) to the inferred *Oct4-Sox2* motif consensus ATT<u>TG</u>CATAA<u>C</u>AAT<u>G</u>, two in the motif’s core ATTT<u>G</u>CATAA<u>C</u>AATG. As expected, the contribution is low at the mismatches. See Figure S1H for the predicted binding contributions. **H)** Experimentally observed ATAC-seq profiles across the *Btbd11* enhancer for the wildtype (solid line) and the CRISPR mutated sequence (dotted line) shows that the low-affinity *Oct4-Sox2* motif is necessary for the full wildtype accessibility across the region.

The discovered motifs showed that both models learned similar *Oct4-Sox2, Sox2, Klf4* and *Zic3* motifs (Figures 1C and S1B-C), consistent with experimental findings that Oct4, Sox2 and Klf4 have a pioneering role in mESCs.^55–57^ The *Nanog* motif was only learned in the TF binding model but not by the accessibility model (Figure 1C). This suggests that Nanog is not a pioneer TF, consistent with Nanog’s dependency on other TFs for binding^36^ and the relative lack of observed chromatin accessibility effects upon its degradation.^58^

We next mapped all individual motif instances within the 1 kb genomic regions, seeking to map as many contributing low-affinity motifs as possible while maintaining confidence that they are specific for the assigned TF (Methods). Briefly, we performed CWM scanning^36^ using the contribution scores to ensure that individual mapped motifs were learned by the model and Jaccardian similarity with the CWM to identify the motif. The returned motif mappings go below typical PWM consensus matches (<85%),^59,60^ thus possess a broader distribution of sequence affinities than those from traditional PWM scanning (Figures 1D and S1D, Methods), yet are still bound in *in vitro* protein-binding experiments^61–64^ (Figure S1E). For simplicity, we refer to an imperfect motif match as “low-affinity”,^2^ regardless of its *in vitro* binding affinity.

To validate that this approach correctly mapped functional motifs, we chose the composite *Oct4-Sox2* motif as an example because longer motifs show a larger number of mismatches to the consensus sequence while still being functional.^35,65^ We found that the *Oct4-Sox2* motifs mapped by the accessibility model consistently showed sharp footprints by the corresponding TF in the ChIP-nexus data, although these binding data were never seen by the accessibility model (Figures 1D and 1E). Furthermore, the motifs with either high, medium or low PWM scores showed decreasing average footprint heights for Oct4 and Sox2, consistent with *in vivo* TF occupancy broadly mirroring motif affinities.^26,66^ Interestingly, the weaker *Oct4-Sox2* motifs showed a full-length *Oct4* component with an almost non-existent *Sox2* motif component in the sequence, but nevertheless showed distinct ChIP-nexus footprints for both Oct4 and Sox2 (arrows in Figure 1E), suggesting that these motifs are still bound by an Oct4-Sox2 complex. This explains previous observations where strong pioneering effects were observed for regions with an *Oct4* motif in mESCs.^22,67^ In summary, the vast majority of mapped motifs, including those with lower PWM scores, showed independent experimental evidence for being functional.

As a comparison, we mapped motifs in the same genomic regions using traditional PWM scanning^59,60^ (Methods). A low match score requirement yielded much higher numbers of mapped motifs, the majority of which was not mapped with our method (Figure S1G). While this is expected for low PWM scores, almost no specific binding was observed even among the top 5,000 motifs with the highest PWM scores if they were not also mapped with our approach (Figure 1F). This shows that PWM scanning produces false positives even at higher match scores and that deep learning models are more accurate in mapping functional motifs, consistent with previous results.^35,36^ We note that once individual motif instances are mapped and identified through deep learning models, PWM scores still serve as a robust independent metric to measure motif strength (Figure S1E, Methods).

Finally, we performed a CRISPR/Cas9-mediated genome editing experiment to test whether low-affinity motifs indeed contribute to chromatin accessibility in a detectable manner. We selected the previously examined intronic *Btbd11* enhancer^36^ since we discovered that it has a low-affinity *Oct4-Sox2* pioneer motif with key mismatches across both the *Oct4* and *Sox2* motif components (Figure 1G). We performed two point mutations that were predicted to abolish the motif’s pioneering effect (Figures 1G and S1H, Methods) and performed ATAC-seq experiments on wildtype and mutated cells (Figures S1I and S1J). While the observed effect differed slightly from the predicted effect (Figure S1K), the accessibility in the region around the mutated motif was significantly decreased, showing that the low-affinity *Oct4-Sox2* motif has a measurable effect on chromatin accessibility (Figure 1H). Taken together, these results suggest that we accurately mapped low-affinity motifs that contribute to chromatin accessibility.

### Low-affinity pioneer motifs are contextually more enhanced

We next set out to analyze how much the motif of each TF contributes to chromatin accessibility and how much this depends on the motif’s intrinsic binding affinity versus its surrounding genomic context. To do so, we extracted two measurements from our models across all mapped motifs, isolation scores and context scores, which we will use throughout this work.

Isolation scores (also called marginalization scores^21,68^ or global importance scores^69^) measure the predicted effect of each motif sequence *in isolation*, when injected into controlled, randomized sequences (Figure 2A, log-fold-change over control, Methods). For a TF binding model, the isolation scores for mapped motifs have been shown to be a proxy for relative motif affinities.^68^ In contrast, context scores, a motif-centric expansion of *in silico* mutagenesis^70,71^ or the necessity test,^72^ measure the predicted loss when a motif is perturbed in its genomic sequence context^36^ (Figure 2B, Methods).

**Figure 2.**
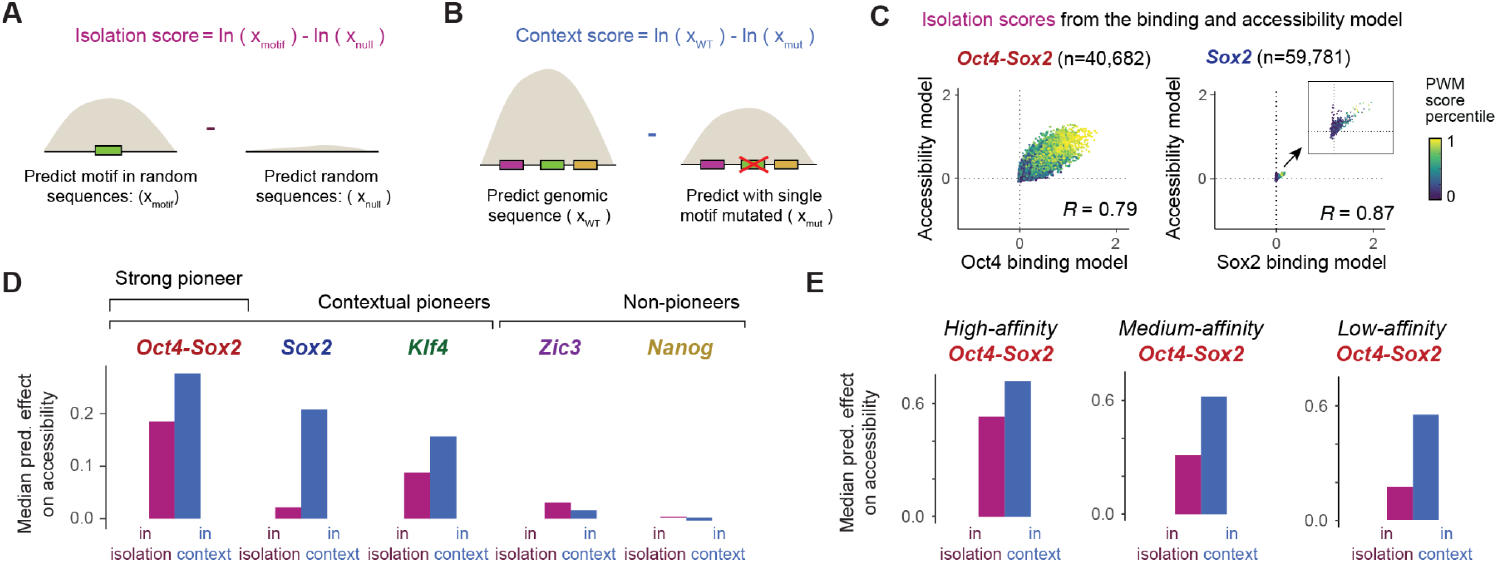
Low-affinity motifs can act as contextual pioneers. **A)** Isolation scores from binding and accessibility models were obtained by injecting each uniquely identified motif sequence across 256 random *in silico* sequences, averaging the total reads predicted across injected and uninjected sequence trials and measuring the log-fold-change of the injected averaged sequence effects vs uninjected averaged sequence effects. **B)** Context scores from binding and accessibility models were obtained by mutating each mapped motif coordinate with 16 random sequence replacements, averaging the total reads predicted across mutated sequence trials and measuring the log-fold-change effects of the wildtype sequence effects over the averaged mutation effects. **C)** Comparison of the isolation scores obtained from the binding model (x-axis) and the accessibility model (y-axis) for the *Oct4-Sox2* (left) or the *Sox2* (right) mapped motifs, colored by their PWM score percentiles, showing that the isolation scores of both models correlate with PWM scores. **D)** For each set of motifs mapped by binding and accessibility models, the median isolation score for each motif(left) shows its relative pioneering strength, while the median context scores (right) show the motif’s impact in the genomic context. This reveals *Oct4-Sox2* as the strongest pioneer motif and the contextual effects for *Sox2* and *Klf4*. **E)** For *Oct4-Sox2* motifs motifs mapped by only accessibility models, median isolation and context scores predicted by the accessibility model of high, medium and low affinity motifs (5k each, based on PWM score) show that low-affinity motifs receive a bigger boost in context than high-affinity motifs.

Intuitively, a motif’s isolation and context score should be similar if the motif’s contribution is independent of other sequence features in the genomic context. If, however, motifs have bigger context scores than isolation scores, it suggests that they are highly contextual and may function cooperatively with other motifs. The advantage of this approach is that it directly represents a measurement of a motif’s entire effect within a genomic context, without having to perform a more focused cooperativity analysis for each motif pair, which we will conduct later.

To validate these scores, we first asked whether the isolation scores from the TF binding model relate to motif affinities and whether the accessibility model also learned affinity proxies, as previously observed.^21^ We found that indeed for both models, the isolation scores for each motif were correlated with PWM scores (Figures 2C and S2A) and in vitro protein-binding microarray experiments^61–64^ (Figure S1F and Table S3, Methods). Furthermore, the isolation scores were highly correlated between the two models (Figures 2C, S2A and Table S3, Spearman correlation 0.67-0.87). This is surprising since the models were trained independently on different types of experimental data and the context scores for each motif were only moderately correlated (Figure S2B and Table S3). This suggests that relative motif affinities are indeed represented by the isolation scores and that they were learned independently by both models.

Having obtained some validation, we next asked how the TF motifs differed from each other in their pioneering strength and dependency on genomic context. Since the TF motifs were learned together in the accessibility model, each TF motif’s average isolation score should represent its intrinsic baseline contribution towards pioneering. To have a broad representation of all motifs, including those of non-pioneer TFs, we included motifs learned by both models. This revealed that *Oct4-Sox2* motifs possessed the strongest pioneering effects in isolation, with a relatively small boost in context (Figure 2D). This is consistent with *Oct4-Sox2’*s inherent cooperativity from Oct4 and Sox2 binding, the motif’s strong influence on the binding of other TFs^36,73^ and the strong loss of accessibility upon Oct4 depletion.^58,67,74^ On the other end of the spectrum, *Zic3* and *Nanog* motifs produced poor correlations between binding and accessibility effects (Spearman correlation 0.21-0.30, Table S3), suggesting that they do not pioneer chromatin in these cells. Interestingly, *Sox2* and *Klf4* motifs had weak isolation scores but strong context scores. For example, the *Sox2* motif showed on average over 10-fold stronger pioneering effects when placed in a genomic context. This suggests that the *Sox2* and *Klf4* motifs benefit more from the surrounding genomic context.

The strong contextual effects of *Sox2* and *Klf4* raise the question whether these are particularly cooperative pioneer TFs, or alternatively, whether weak pioneer motifs in general tend to get bigger contextual boosts. If the latter is true, we would expect to see the same effect even among motifs for the same TF, i.e. low-affinity motifs should get boosted proportionally more in their genomic context than the high-affinity counterparts. We therefore analyzed the set of *Oct4-Sox2* motifs of high, medium and low affinity from the accessibility model as described earlier and compared the average context and isolation scores in each group (Figure 2E). We found that the high-affinity motifs had a ∼35% greater effect in context than in isolation, while the low-affinity motifs had a ∼300% (3-fold) greater effect. Likewise, the *Sox2* and *Klf4* motifs showed lower isolation scores with lower affinities, but the context scores decreased less, showing their increased reliance on context for their effects (Figures S2C and 2D). These results suggest a general tendency of motifs with weaker effects to be more enhanced in a genomic context, consistent with low-affinity motif mutations being able to cause strong phenotypic effects.^5–7^ This points to a need to understand the sequence rules and mechanisms underlying this contextual enhancement.

### Identical motif sequences have different effects depending on adjacent pioneer motifs

To understand these contextual effects, we tested whether motifs are enhanced by being near other motifs or by being more centrally located within the accessible region, as previously suggested.^19,21,27,37,38,75^ Using the context scores for a motif’s predicted pioneering strength, we asked to what extent these scores are determined by motif proximity, centrality within the ATAC-seq region, or motif affinity itself (Figure 3A). We found that high pioneering effects of a weaker pioneering motif were best predicted by small distances to the strongest pioneer motif in the region (Spearman correlation 0.37 to 0.38), even more than motif affinity (Spearman correlation 0.06 to 0.11) (Figure 3B and Table S4).

**Figure 3.**
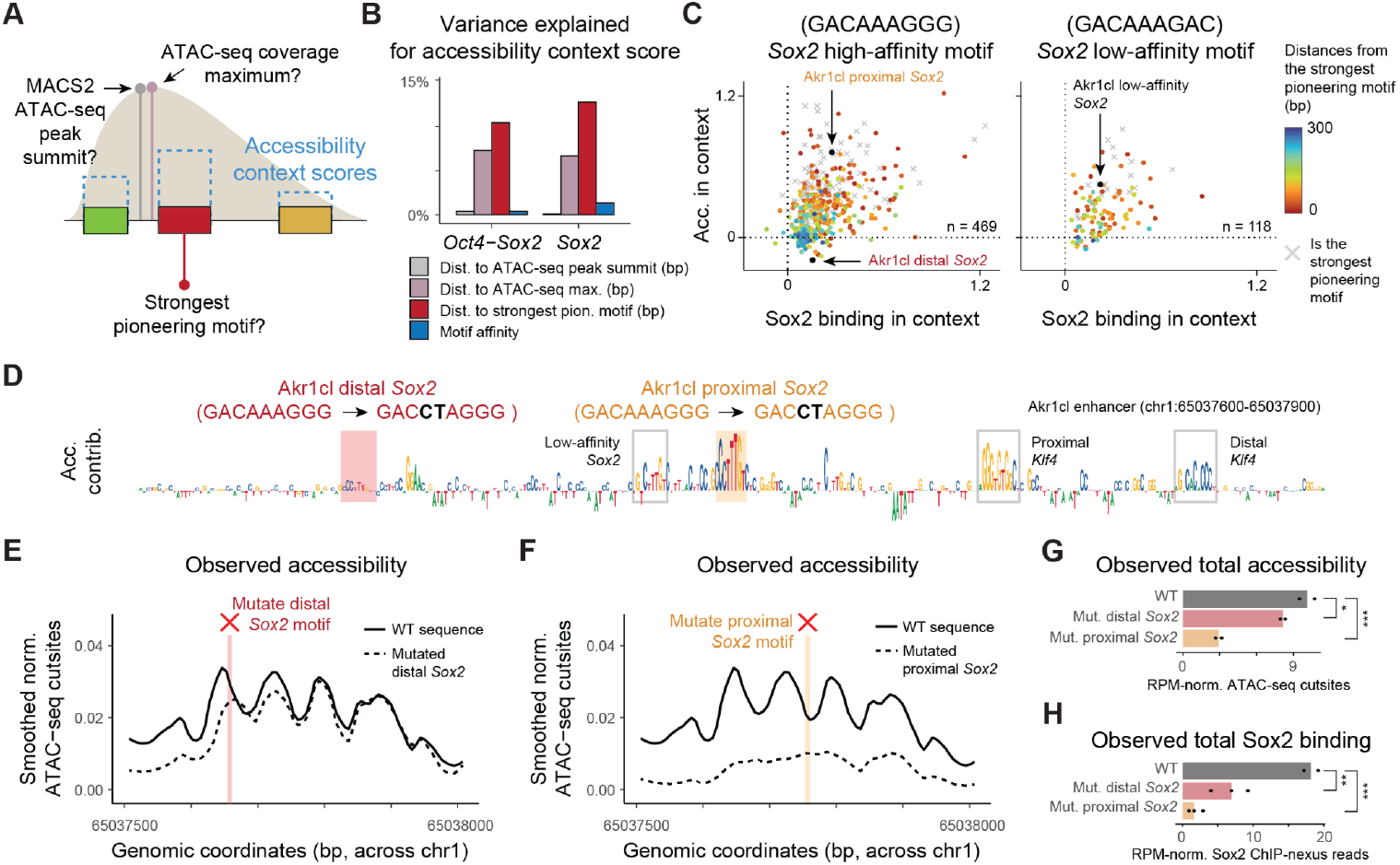
Distance-dependent arrangements of pioneer motifs drive strong effects. **A)** Graphic depicting different measurements across every mapped motif as follows: (1) the distance from the center of the considered motif to the center of the strongest pioneer motif (defined as possessing the highest accessibility context score), (2) the distance from the accessibility peak summit via MACS2 and (3) the distance from the ATAC-seq fragment coverage maximum. **B)** Barplots measuring the variance explained from each measurement to explain *Oct4-Sox2* and *Sox2* accessibility contribution scores using independent linear regression. Strongest pioneering motifs were excluded. **C)** Binding and accessibility context scores of mapped *Sox2* motifs which contain precise sequence matches to a high-affinity *Sox2* (GACAAAGGG) or a low-affinity *Sox2* (GACAAAGAC) motifs, colored by the distance of each motif to the strongest pioneer motif in the region. Gray crosses indicate the strongest pioneer motifs. **D)** Accessibility contribution scores of the Akr1cl upstream enhancer (chr1: 65037600-65037900) with contributing motifs marked and the two sequence ubstitutions of the distal and proximal *Sox2* sequences annotated. **E-F)** Smoothed experimental wildtype (solid line) and CRISPR mutant (dashed line) ATAC-seq cut site profiles across the *Akr1cl* enhancer compare the effects of **(E)** mutating the distal *Sox2* motif and **(F)** mutating the proximal *Sox2* motif. **G-H)** After RPM-normalization, median (colored bars) total **(G)** ATAC-seq cut sites or **(H)** Sox2 ChIP-nexus reads across replicates (black dots) occurring across chr1: 65037600-65037900 across wildtype (black), distal *Sox2* mutant (red) and proximal *Sox2* mutant (orange) experiments. Significance was tested via a one-tailed t-test between replicate experiments (black dots).

This suggests that the same motif may mediate substantially stronger pioneering effects when closer to a strong pioneer motif. To visualize this correlation, we plotted the context scores for all motifs of the same sequence, either a high-affinity *Sox2* (GACAAAGGG) or a low-affinity *Sox2* (GACAAAGAC), and colored them by the distance to the strongest pioneer motif (Figure 3C and Table S4). This showed a strong distance correlation (Spearman correlation 0.28 to 0.47, Table S4) and illustrated how even a low-affinity *Sox2* can have strong effects when nearby a strong pioneer motif.

To experimentally test whether identical motif sequences have different effects on chromatin accessibility, we performed CRISPR/Cas9 editing on the *Akr1cl* enhancer. This region has two high-affinity *Sox2* motifs of the exact same sequence but with different accessibility context scores. Both are bound by Sox2 (Figures S3A and S3B), but one is located proximally to a low-affinity *Sox2* motif and two *Klf4* motifs, while the other is positioned more distally (Figure 3D). After selecting successful CRISPR/Cas9 clones with targeted point mutations in each motif, we performed ATAC-seq as well as Sox2 ChIP-nexus experiments to confirm the loss of Sox2 binding (Figures S3C-E).

Mutating the distal *Sox2* motif had a slight, yet significant decrease in accessibility levels (Figures 3E and 3G), but mutating the proximal *Sox2* motif produced more than a three-fold reduction in accessibility (Figures 3F and 3G). These results differed slightly from the predicted effects (Figure S3F), but confirmed that mutating the proximal *Sox2* motif showed a stronger reduction in accessibility than mutating the distal motif, despite both motifs and their mutations being identical in sequence.

Finally, we investigated whether the differential effect on accessibility also affected Sox2 binding. We found that Sox2 was still bound to the intact proximal motif when the distal Sox2 motif was mutated (Figure S3G). However, when the proximal Sox2 motif was mutated and chromatin accessibility was strongly reduced, the intact distal Sox2 motif was no longer bound by Sox2 (Figure 3H and S3H). This suggests a hierarchy of TF binding where binding to strong pioneer motifs promotes binding to motifs with weaker pioneering effects in the same region.

### Pioneering cooperativity occurs through nucleosome-range soft syntax

Since the effects of motifs strongly depended on adjacent motifs, we systematically queried the model to identify the rules by which two motifs cooperate. We performed *in silico* perturbations for each motif pair by mutating each motif alone or together within its genomic context. We then tested whether the joint effects (WT over the double mutant) are more than the sum of each motif’s effect (each single mutant over the double mutant)^19,37,76^ (Figure 4A).

**Figure 4.**
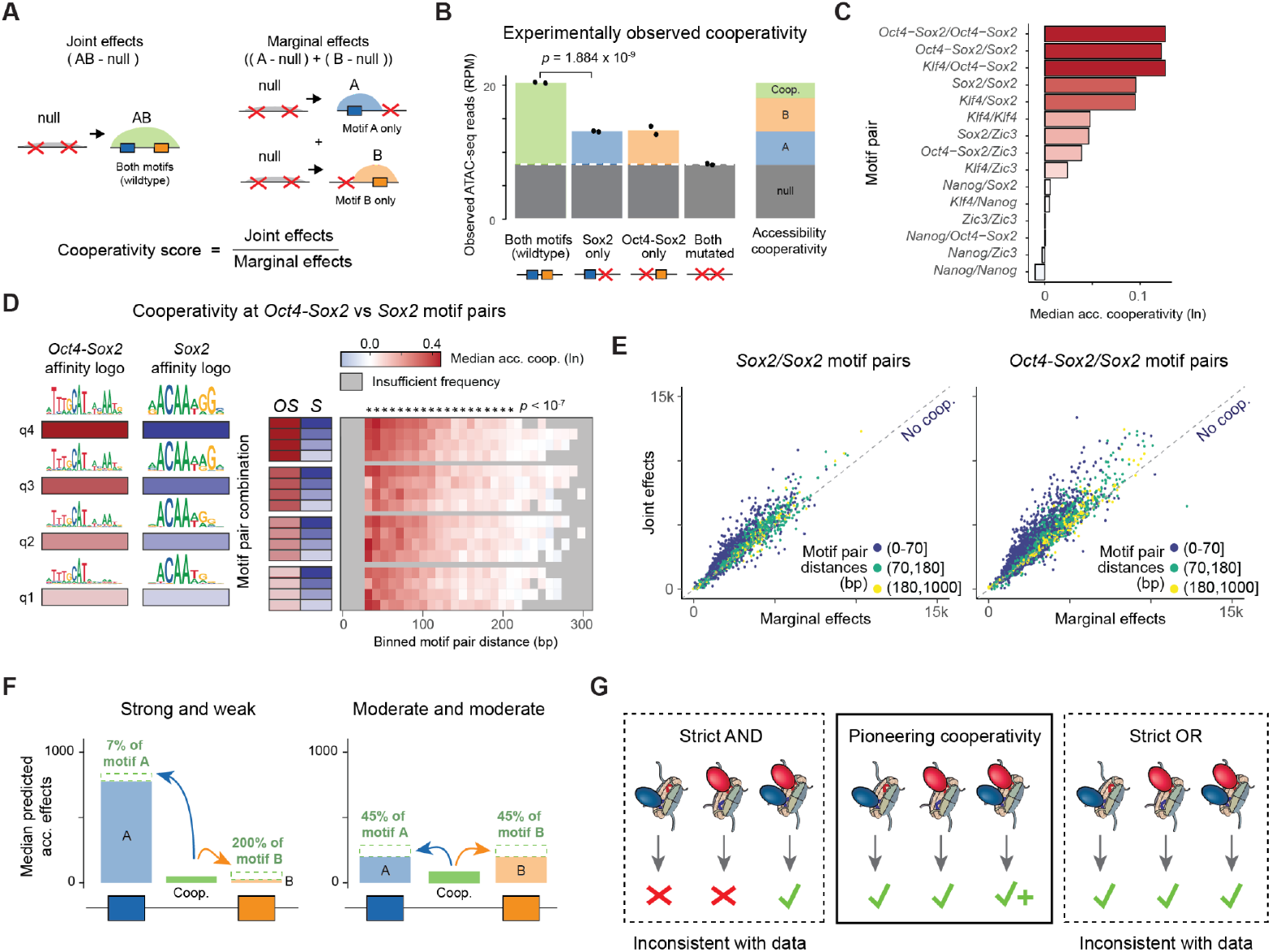
High-affinity and low-affinity motifs cooperate within nucleosome distances to increase accessibility. **A)** Graphic defining superadditive accessibility cooperativity. **B)** Across the Btbd11 intronic enhancer, CRISPR mutations were conducted to establish cell lines representing the four motif pair configurations mutating the *Sox2* and low-affinity *Oct4-Sox2* motif. Next, ATAC-seq experiments were performed and RPM-normalized reads were collected across chr10:85539400-85539800 from the wildtype and each CRISPR experiment. Joint effects and marginal effects of each motif were calculated across each replicate set, with significance derived from a one-tailed t-test. To the right, calculated cooperative effects (green) are stacked upon individual observed effects of motifs. **C)** Median accessibility cooperativity across all motif pairs of key mESC motifs occurring within 300 bp of one another, also colored by median accessibility cooperativity. **D)** Median accessibility cooperativity between *Oct4-Sox2*/*Sox2* motif pairs across quartiles of motif affinity and motif pair distances, binned to 10 bp. Motif affinity and binned distance configurations with fewer than 20 motif pairs were not considered (gray boxes). Cooperativity significance was derived from a one-tailed Wilcoxson test comparing whether *Oct4-Sox2*/*Sox2* motif pairs arranged at select binned distances were more cooperative than *Oct4-Sox2*/*Sox2* motif pairs arranged at very long distances (>400 bp) with multiple-comparison Bonferroni corrections and an adjusted significance cutoff of *p* < 10^-7^. **E)** Comparison of joint and marginal predicted accessibility effects for the *Sox2*/*Sox2* (left) and *Oct4-Sox2*/*Sox2* (right) motif pairs, colored by the pairs’ relative center-to-center distances to one another. Motif pairs with higher joint effects than marginal effects were interpreted as cooperative. **F)** As denoted in **(A)**, median accessibility marginal effects (blue and orange) and calculated cooperativity levels (green) of all pioneer motif pairs containing *Oct4-Sox2, Sox2* and *Klf4* arranged within 200 bp of one another. The left panel depicts motifs with the strongest 20% predicted accessibility context scores paired with motifs with the weakest, non-zero 20% predicted context scores. The right panel depicts paired motifs both within the middle 20% of accessibility context scores. **G)** Implications for the mechanisms of nucleosome-mediated cooperativity.

Using this analytical approach, we confirmed cooperativity at the *Btbd11* enhancer where we mutated either the low-affinity *Oct4-Sox2* motif, the *Sox2* motif, or both motifs, using CRISPR-editing and measured the changes in ATAC-seq Finally, we investigated whether the differential effect on accessibility also affected Sox2 binding. We found that Sox2 was still bound to the intact proximal motif when the distal *Sox2* motif was mutated (Figure S3G). However, when the proximal *Sox2* motif was mutated and chromatin accessibility was strongly reduced, the intact distal *Sox2* motif was no longer bound by Sox2 (Figure 3H and S3H). This suggests a hierarchy of TF binding where binding to strong pioneer motifs promotes binding to motifs with weaker pioneering effects in the same region. accessibility on the validated clones (Figures S1I-J). While the observed effects differed slightly from the predicted effects, the joint effect of both motifs was larger than the added effects of each motif, confirming cooperativity (Figures 4B and S1K).

We then calculated the *in silico* cooperativity scores for all motif pairs across the genome. We found that all pairs of pioneer motifs showed cooperativity substantially above 1 (or above 0 in log-space) (Figures 4C and S4A-B), while the cooperativity was weakened or non-existent when one motif was a non-pioneer motif, such as *Nanog* (Figure S4C). When cooperativity occurred, the cooperativity scores showed distance-dependent enhancements that were similar between different motif pairs and across various motif affinities (Figures 4D-E and S4A-B). Significant cooperativity above that of very distant motif pairs typically occurred within distances of 200 bp and was highest for motif distances of <70 bp (Figures 4D-E and S4A-B), thus a soft syntax that evokes a nucleosome-mediated mechanism.^36^

The rules of cooperativity are consistent with our initial observation that weak pioneer motifs tend to have strong effects in their genomic context. When weak motifs cooperate with stronger motifs, the resulting accessibility increase is relatively higher for the weak motif than the strong motif, while two motifs of similar strength benefit from cooperating more equally (Figure 4F). For low-affinity motifs, their average contextual enhancement is further enhanced by the fact that weak motifs are only functionally relevant (and mapped by our models) when they cooperate with stronger pioneer motifs, while strong pioneer motifs may sometimes function on their own (Figure 4F). Thus, functional motifs of very low affinity are by definition cooperative and strongly contextually enhanced.

Since the strongest effects were observed when two motifs are pioneer motifs and likely found on the same nucleosome, we considered the mechanistic implications (Figure 4G). There is accumulating evidence for nucleosome-mediated cooperativity,^77–80^ but the exact mechanisms are not understood. The involvement of two pioneer TFs suggests that their relationship is not simply a hierarchical one where one pioneer TF enables the binding of another TF to accessible DNA.^78,79^ Instead, the two pioneer TFs likely cooperate in removing the same nucleosome, and our analyses rule out two extreme models for this cooperativity. First, we can exclude a model where two or more pioneer TFs must be present together on the same nucleosome to remove it (strict AND in Figure 4G). Such a strict requirement for multiple TFs is inconsistent with the model’s prediction that each pioneer TF produces a certain amount of accessibility per motif. Second, we can exclude a model where each pioneer TF independently removes the same nucleosome with a certain probability, causing increased accessibility across the cell population (strict OR in Figure 4G). If this were the case, this would produce less than additive effects (Figure S4D). This suggests that two pioneer TFs quantitatively enhance each other’s effects in removing a nucleosome, and that this mechanism applies to low-affinity motifs.

### Modeling suggests that pioneering cooperativity changes the maximum regulatory potential

We next explored a kinetic modeling framework to understand the implications of nucleosome-mediated cooperativity, especially in response to changing TF concentrations. Based on thermodynamics, increasing TF concentrations lead to increased binding of the TF to a motif, roughly linearly at intermediate TF levels and then plateauing towards 100% at high concentrations. 10,81,82 For low-affinity motifs, higher TF concentrations are required to reach full TF occupancy. 17 How motifs and their cooperative effects modulate chromatin accessibility as a function of TF concentration is not known.

Our approach is similar to previous thermodynamic equilibrium models in that it considers TF binding to a closed and an open conformation for which the TF has different affinities 27,77,79 (Figure 5A). While it can be parameterized to correspond to a system at thermodynamic equilibrium, it includes the option of modeling the TF-nucleosome dynamics away from equilibrium to account for the irreversible nature of nucleosome removal by ATP-dependent remodelers 83,84 (Figures S5A and S5B). In this kinetic model, the strength of pioneering at a given TF concentration depends on the motif’s affinity (1/g), the nucleosome intrinsic affinity for DNA (k_close_/k_open_), the nucleosome’s effect on destabilizing TF binding (β) and the TF’s ability to increase the rate of opening through nucleosome remodeling (o) and possibly preventing nucleosome reformation (c) (Figures 5A and S5C, Methods).

**Figure 5.**
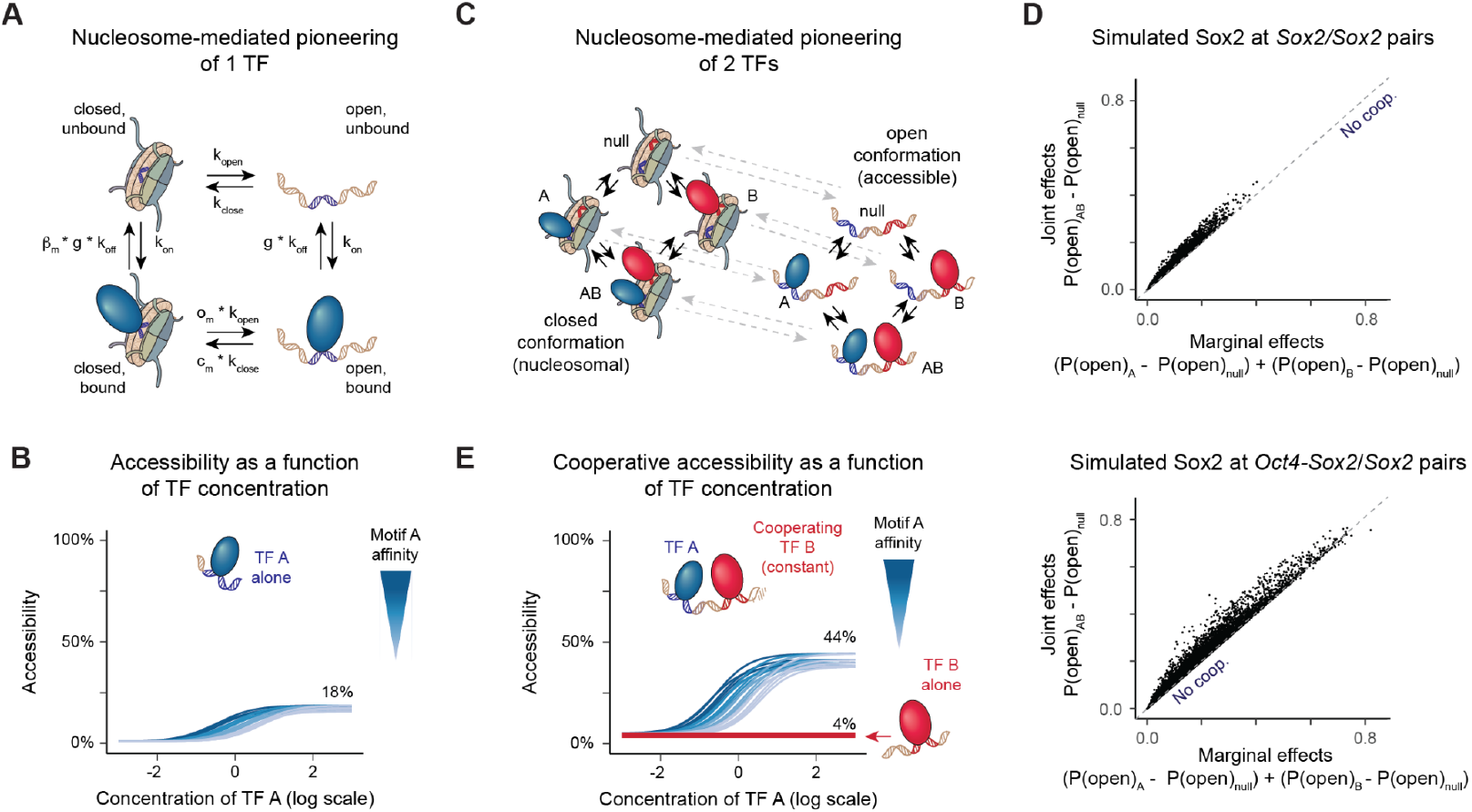
Pioneering cooperativity is predicted to change the TF dose response curve. **A)** Graphic depicting a kinetic model out of equilibrium of bound TF effects on nucleosome removal and reformation. Parameter definitions are as follows: affinity (1/g, factor decrease in unbinding rate on the motif), TF association rate (k_on_), TF disassociation rate (k_off_), chromatin opening rate in absence of bound TF (k_open_), chromatin closing rate in absence of bound TF (k_close_), nucleosomal effect upon TF disassociation rate (β_m_ when motif is present, β_n_ when motif is absent), TF effect on nucleosome removal (o), TF effect on nucleosome reformation (c). **B)** 1 TF kinetic model simulations (n=5) measuring the accessibility state, measured as probability towards the open conformation, upon pioneer TF Sox2 binding over a range of motif affinities across changing concentrations. Parameter ranges can be found in Table S6. **C)** Graphic depicting an expansion of the kinetic model shown in **(A)**, allowing for simultaneous effects of two pioneer TFs. **D)** 2 TF kinetic model simulations (n=50,000) comparing joint and marginal accessibility states, measured as probability towards the open conformation, for Sox2 binding at a *Sox2/Sox2* motif pair (top) and an *Oct4-Sox2/Sox2* motif pair (bottom). Parameter ranges can be found in Table S7. Simulations that returned higher joint effects than marginal effects were interpreted as cooperative. **E)** 2 TF kinetic model simulations (n=5) measuring the accessibility state, measured as probability towards the open conformation, upon two pioneer TFs binding over a range of TF A motif affinities across changing concentrations of TF A. TF A was modeled from Sox2 binding dynamics *in vivo* with the same parameters as **(B)**. TF B was modeled with the same pioneering parameters, but with static motif affinity and TF concentration, such that its unchanging effect on accessibility in the corresponding 1 TF kinetic model is depicted with a red line. Though TF B affinity and concentration was kept constant, the presence of TF B caused accessibility states to be cooperatively enhanced beyond those of TF A alone **(B)**. Parameter ranges can be found in Table S7.

We applied this kinetics model to Sox2 since we could choose parameter ranges in agreement with measured Sox2 concentrations in mESCs^85^ and Sox2 binding dynamics *in vitro* and *in vivo*.^86–88^ By modeling a single *Sox2* motif and choosing diverse parameter sets representing variability of individual genomic regions within plausible ranges (Methods), we observed Sox2 pioneering strength proportional to motif affinity (Figure S5D), as observed in the accessibility deep learning models (Figure 2C).

We then explored how chromatin accessibility depends on the motif affinity and Sox2 concentration. With rising Sox2 concentrations, the levels of accessibility increased almost linearly. Lower motif affinities shifted the curve towards higher Sox2 concentrations, but the curves eventually reached the same plateau of maximum accessibility, which was typically less than 100% (Figure 5B). Single-molecule footprinting studies indeed show that typically only a fraction of nucleosomes are removed at accessible regions, despite *in vivo* enhancers often containing multiple pioneer motifs.^78,79,89^ Our modeling suggests however that this may not be because the TF concentrations are too low, but rather that there is a maximum fraction that can be achieved with those TFs. For a single TF, we see that this maximum depends on the nucleosome’s affinity for DNA and the ability of the TF to promote the open conformation (Figures 5B and S5C).

We then asked how this dose response curve might change when Sox2 cooperates with another TF. We expanded the kinetics model to simultaneously consider two TFs binding to their corresponding motifs (Figure 5C, Methods). We assumed that when both TFs are bound (AB in Figure 5C), the factors by which each TF increases the opening rate (o_m_) multiply, which can be interpreted to correspond to an additive effect on lowering the activation energy of the transition (Methods). Under this assumption, we could readily simulate the cooperativity effects predicted by our accessibility deep learning model of Sox2 binding to *Sox2*/*Sox2* and *Oct4-Sox2*/*Sox2* motif pairs (Figure 4E) over a range of biologically plausible motif affinities and parameter settings (Figure 5D, Methods).

To understand how pioneering cooperativity affects the concentration curve, we then simulated a simpler scenario where Sox2 was again binding to a *Sox2* motif, but this time in the presence of a cooperating motif whose pioneer TF was kept at a constant low concentration (Methods). With rising Sox2 concentrations, the total fraction of accessible chromatin increased more rapidly (Figure S5E) and reached a higher plateau compared to that of the single motif (Figure 5E), suggesting that pioneering cooperativity extends the regulatory potential of the motif. This effect was distinct from the impact of motif affinity, which did not change the maximum accessibility state (Figures 5B and 5E). Thus, the modeling suggests that there is a maximum state of accessibility when a TF achieves full occupancy on nucleosomal and naked DNA (Figure S5F). This maximum impact is set by the pioneering strength of the TF and the cooperative motif arrangement in the genomic region (Figures 5E and S5G). Thus, cooperative motif arrangements have a higher accessibility impact and respond faster to concentration changes.

This raises the possibility that the widespread impact of low-affinity motifs in developmental enhancers may not be due to the effects of motif affinity *per se*, but due to the enhancer properties conferred by the cooperativity that is inherent to low-affinity motifs. While low-affinity motifs do not cooperate more strongly than high-affinity motifs (Figures 4D and S5G), they are abundant and arise more easily during evolution, and when their effect is detectable, they are by definition in a cooperative arrangement.

### Cooperating motifs are more sensitive to changing Oct4 concentrations *in vivo*

We next tested the modeling predictions *in vivo* by analyzing an accessibility time-course data set in mESCs where the concentration of Oct4 gradually decreases after doxycycline withdrawal (0h, 3h, 6h, 9h, 12h and 15h)^67^ (Figure S6A). To test whether loss of Oct4 was the main driver for the observed accessibility changes, we trained separate models for each time point and analyzed the motifs as before (Figure 6A and Table S2). We found that the contributions of *Oct4-Sox2* motifs decreased over time, while other motifs maintained similar contributions, including *Sox2* and *CTCF* (Figures 6B and S6B). The *Oct4-Sox2* motif was still discovered by the model at 12h, when Oct4 was at 7% of its original concentration, but not at 15h, when Oct4 was at 2.5% and the *Oct4-Sox2* motif contributions were minimal. These results confirm the robustness of the model training and interpretation (Figures S6C and S6D) and suggest that the decreasing contribution of the *Oct4-Sox2* motif is the main driver for the observed accessibility changes.

**Figure 6.**
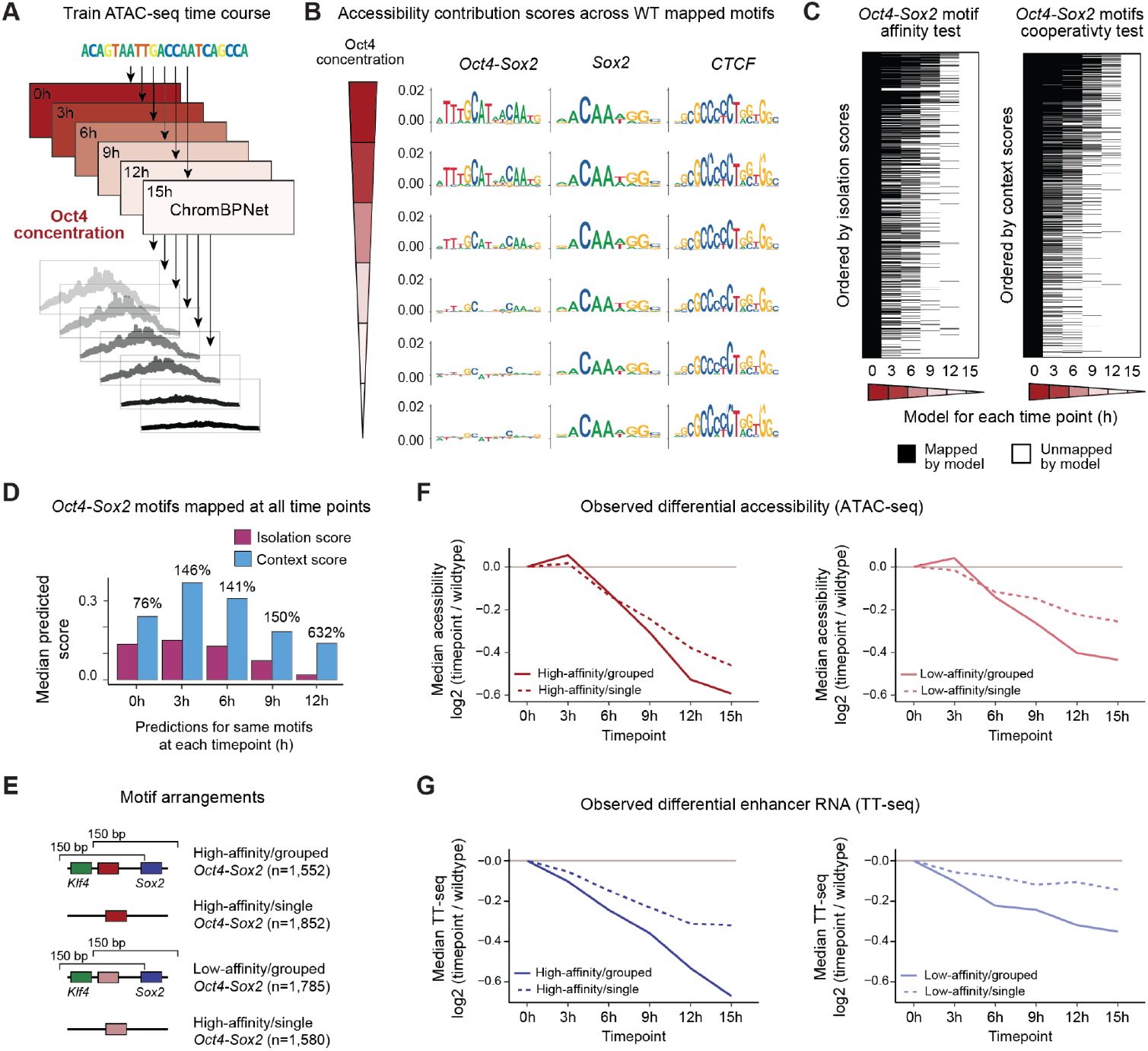
Cooperative motif arrangements bolster motif pioneering across changing TF concentrations. **A)** ChromBPNet accessibility models were trained on ATAC-seq data collected over decreasing Oct4 concentrations via doxycycline withdrawal experiments. All models were provided the original set of 152,827 regions for training, tuning and testing. **B)** Accessibility CWMs measured across all Oct4 concentration timepoints using mapped motifs derived from the R1 mESC accessibility model (Figures 1B and 1C) for *Oct4-Sox2, Sox2* and *Ctcf* motifs. **C)** Heatmaps marking which *Oct4-Sox2* motifs mapped by the 0h timepoint model were mapped by the corresponding time course accessibility models, ordered by the motif isolation scores as proxy for motif affinity (left) or the context scores (right), which also incorporate the cooperativity found at these motifs. Both sets of scores were from the 0h timepoint model. **D)** As *Oct4-Sox2* motifs remained mapped throughout the time course (x-axis), we measured median context and isolation scores with reported ratios between each median context and isolation score. **E)** Graphic denoting arrangements of single/grouped and strong/weak *Oct4-Sox2* motifs. *Oct4-Sox2* motifs were derived from the 0h timepoint model, while high- and low-affinity motifs were classified here as the top and bottom tertiles of measured PWM scores. **F)** Median experimental differential accessibility levels measured over the concentration time course across regions with either single (solid line) or grouped (dashed line) *Oct4-Sox2* motifs, subplots depicting high (left) and low (right) affinity groups. Note that there is an initial increase, consistent with Xiong et al. 2022, perhaps because Sox2 gets released from the Oct4-Sox2 complex and binds to *Sox2* motifs. **G)** Median experimental differential enhancer expression levels measured over the concentration time course using TT-seq across regions with either single (solid line) or grouped (dashed line) *Oct4-Sox2* motifs, subplots depicting high (left) and low (right) affinity groups.

We next asked what sequence features best predicted when individual *Oct4-Sox2* motifs were no longer detected and mapped. Based on thermodynamics, we expect that high-affinity motifs are occupied the longest, and thus the *Oct4-Sox2* isolation scores should be a good predictor for the continued motif detection. Our modeling suggested however that the loss of accessibility depends on the *Oct4-Sox2* motif in its cooperative context, and thus the context scores should also be a good predictor. Indeed, we found that while both measurements were predictive, the context scores were the best predictor (Figures 6C and S6E-F), implying that motifs are detected at lower TF concentrations if they are contextually enhanced.

We therefore examined how *Oct4-Sox2* motifs that were still mapped until 12h benefited from their cooperativity with other motifs. In these regions, the only change is that the *Oct4-Sox2* motifs have lower effects in isolation due to the decreasing Oct4 concentrations, while cooperating TFs, such as Sox2, remain present. In this controlled setting, the context scores decreased over time, but they remained higher relative to the decreasing isolation scores. While these motifs received a ∼76% context boost at 0h, the boost was ∼632% at 12h when these motifs were intrinsically much weaker (Figure 6D). This corroborates our conclusion that weak pioneer motifs functionally benefit from their genomic context.

Collectively, these results suggest that an enhancer’s response to changing TF concentrations depends on its cooperative motif arrangements. Specifically, our kinetic model suggested that cooperative arrangements have higher accessibility potential and thus should start with higher accessibility levels, with a faster loss over time. To test this, we sampled regions with exactly one high-affinity, or alternatively one low-affinity *Oct4-Sox2* motif, which were either found in a “grouped” cooperative arrangement with *Klf4* and *Sox2* motifs or had a “single” *Oct4-Sox2* motif (Figures 6E and 6F), while considering confounding variables (Figures S6G-I). For each time point, the loss of accessibility compared to the starting point (0h) was quantified using DESeq2.^90^ Regions with a *Sox2* motif but no *Oct4-Sox2* motif were used as control (Figure S6J).

This showed that grouped *Oct4-Sox2* motifs had a more rapid loss of accessibility over time than single *Oct4-Sox2* motifs (Figure 6F). While high-affinity motifs showed more accessibility loss than low-affinity motifs, both showed bigger effects in the grouped arrangement than when alone. These results corroborate that cooperativity is a critical factor in making an enhancer more sensitive to changing TF concentrations.

Finally, we analyzed whether these regions also show differences in enhancer activity over the same Oct4 depletion time course, measured as newly transcribed enhancer RNA with TT-seq.^67^ This revealed that the levels of enhancer transcription also dropped faster in the regions with grouped rather than single *Oct4-Sox2* motifs (Figure 6G). This mirrors the trends in the accessibility data, consistent with chromatin accessibility and activation by Oct4 being rate-limiting steps at these enhancers. As a control, grouped and single *Sox2* motifs not near an *Oct4-Sox2* motif show no relationship with differential enhancer transcription (Figure S6J). This suggests that the cooperativity observed for chromatin accessibility in response to changing TF concentrations also affects the activity of enhancers, supporting the *in vivo* relevance of this phenomenon.

## Discussion

Low-affinity motifs are important during development and can impact the phenotypic outcome of mutations.^6,7^ They can be critical for an enhancer’s specificity^4–8^ and may help an enhancer respond to high TF concentration ranges.^9,27,30^ However, mapping functional low-affinity motif instances in the genome has been challenging with traditional PWM-scanning methods and mechanistic studies have mostly been focused on TF binding cooperativity, which occurs at close distances of less than ∼30 bp.^6,15,16,30^ The mechanisms by which low-affinity motifs might regulate enhancer function at the level of chromatin accessibility have remained a gap in knowledge.

Here we addressed this gap by training and interpreting a sequence-to-profile deep learning model on chromatin accessibility. By learning motifs in an inherently combinatorial manner in their genomic context, we show that low-affinity motifs can be accurately mapped, as validated by independent TF binding footprints. Model interpretation then revealed that the pioneering effect of a motif is strongly dependent on its arrangement with other motifs, more than the motif’s affinity. When a weak pioneer motif cooperates with a strong pioneer motif nearby, it strongly benefits from this cooperativity and produces measurable effects on chromatin accessibility, as validated by CRISPR/Cas9-mediated editing. This cooperativity tends to follow a soft syntax that produces chromatin accessibility enhancements that are distance-dependent but occur flexibly when two motifs are located within nucleosome range (∼200 bp).

This has evolutionary implications, i.e. that low-affinity motifs can cooperate during pioneering with less constrained syntax than those for cooperative TF binding and thus can more easily evolve *de novo* close to an existing pioneer motif. The higher statistical likelihood of low-affinity motifs arising, the flexible distance requirement for cooperativity and the potential for strong effects all make it very likely that low-affinity motifs are widespread and broadly shape enhancer function during evolution.

Such a role in evolution is consistent with–and might even explain–the observation that TF motifs undergo high evolutionary turnover.^91^ With multiple motifs contributing to an accessible region, there are less constraints on individual motifs, allowing weaker motifs to evolve and disappear to subtly alter an enhancer’s function. If indeed widespread, this could explain why systematic mutagenesis experiments produce phenotypic changes at sequences outside mapped TF motifs^92,93^ and why novel regulatory elements often arise near already accessible regions.^94,95^

We also obtained evidence for a link between motif cooperativity, maximum accessibility and enhancer activity. While chromatin accessibility is known to be a rate-limiting step for enhancer function, our kinetic model of nucleosome-mediated cooperativity suggested that there is an intrinsic ceiling of a TF’s effect on chromatin accessibility, even at excess TF concentrations. This ceiling is determined by the region’s affinity for a nucleosome (nucleosome barrier) and the TF’s ability to pioneer. Given this limit, it is interesting that motif cooperativity increases the maximum impact of a motif. This makes intuitive sense since cooperativity produces effects more than the sum of each motif and thus a cooperating motif produces higher maximum accessibility. We termed such maximum impact as the regulatory potential that the TF has on the enhancer’s function.

The concept of regulatory potential is relevant when the TF’s concentration increases or decreases during development, as we have shown for a time course experiment of decreasing Oct4 concentrations. A higher regulatory potential due to cooperativity means that the TF’s effect on chromatin accessibility is higher, causing the accessibility to drop more rapidly and linger longer with decreasing TF concentrations. We see this effect also reflected in the levels of enhancer transcription, consistent with chromatin accessibility affecting enhancer activity.

Such a regulatory ceiling set by chromatin accessibility is consistent with previous observations. Imaging studies of the early Drosophila embryo show that motifs for the pioneer TF Zelda increase the responsiveness of enhancers to an NFkB concentration gradient.^96,97^ This response appears to be tied to Zelda’s role in pioneering since cooperativity at the level of NFkB binding does not achieve the same response.^98^ Likewise, a low-affinity *Oct4-Sox2* motif at a *Klf4* enhancer mediates a higher transcriptional output in response to Esrrb and STAT in mESCs.^65^ This suggests that an enhancer’s expression pattern during development is not only shaped by motif affinity, but also by the cooperative motif arrangements during pioneering, allowing it to precisely respond to changing TF concentrations, either temporally or spatially across cells in the embryo.

Finally, the distance-dependent soft syntax we observed for chromatin accessibility has mechanistic implications as it suggests that pioneering and TF binding are intertwined. When interpreting a BPNet model predicting TF binding in mESCs, we found that the TFs often cooperate though soft syntax in a directional manner.^21,35,36,51^ This suggests that TFs enhance each other’s binding through their effect on chromatin accessibility and that there is a hierarchy by which TF binding occurs *in vivo*. TFs with strong pioneer motifs, i.e. Oct4 and Sox2 to the *Oct4-Sox2* motif, likely bind first, while non-pioneer TFs such as Nanog mostly bind when the chromatin is already accessible. Our CRISPR editing experiments at the *Akr1cl* enhancer further support and expand upon this model. The *Sox2* motif found in a context where it produces strong pioneering was necessary for Sox2 to bind to the identical *Sox2* motif with weak pioneering effects, but not vice versa. This suggests that motifs in a region are bound in a hierarchical manner, with strong pioneer motifs becoming bound before weaker ones, even when these are identical motif instances of the same TF.

How exactly TFs cooperate in pioneering remains an open question. Our kinetic model of nucleosome-mediated cooperativity readily simulated our data, consistent with TFs having a direct effect on the rate of ATP-dependent nucleosome removal. We note however that multiple regulatory steps contributed to the simulated opening effects. For example, our kinetically modeled TFs were set to possess slightly more affinity for DNA in the open versus closed state. This parameter is the main determinant of pioneering strength in classically-employed thermodynamic equilibrium models of nucleosome-mediated cooperativity^27,77,79^ (Figure S5B). Such a mechanism contradicts the concept of a pioneer TF that exclusively opens chromatin through its ability to bind nucleosomal DNA,^18^ yet it is a plausible contribution to pioneering. Binding nucleosomal DNA is structurally more challenging than binding naked DNA,^99,100^ and this is true for the Oct4-Sox2 heterodimer binding to the *Oct4-Sox2* motif,^101^ the strongest pioneer motif in mESCs. Furthermore, TF binding in the open state could reduce the rate of nucleosome reformation or promote the removal of neighboring nucleosomes.^84,102^ This could, for example, explain the additional accessibility that is observed when enhancers are active.^21^ Furthermore, given the numerous nucleosome variants and modifications, it seems plausible that a variety of hierarchically layered mechanisms contribute to making chromatin accessible,^100^ which can be addressed in future mechanistic studies.

### Study limitations

Interpretation of sequence-to-function deep learning models to deduce the underlying molecular mechanisms relies on the assumption that these models have been trained on sequences with sufficiently diverse motif affinities and arrangements to correctly learn the sequence rules. Models have been shown to robustly predict motif affinity effects and cooperativity effects,^19,21,68^ but there is a risk that predictions of multiple motif perturbations are out-of-distribution for stable model predictions.^32^ To confirm the inferred syntactic rules, experimental validations were conducted *in vivo* but were limited to a few individual genomic loci. Likewise, the scope of the work is limited to a single cellular context (mESCs) across five key pluripotency TF/motif pairs. While our deep learning models returned additional motifs that contributed to chromatin accessibility in mESCs (Figure S1C), we chose to limit our work to well-studied TFs so we could validate our findings with high-quality ChIP-nexus binding data and literature knowledge. Moreover, our kinetic modeling is limited to two states (closed/open) with direct transitions and does not include partially unwrapped conformations or modified nucleosomes.^103,104^ Finally, we showed that cooperative motif arrangements affect chromatin accessibility and enhancer transcription, but did not examine how this affects gene expression since this is a complex unresolved problem.^105–107^ Future studies with an expanded scope will reveal how broadly applicable our findings are.

## Supplemental files

Supplemental file 1: Excel spreadsheet with Supplementary Tables 1-7 in separate panels. Supplementary Table 1 contains performance metrics and architectures during deep learning model optimization. Supplementary Table 2 contains performance metrics and architectures of finalized deep learning models. Supplementary Table 3 contains correlations of predicted motif effects and affinities.

Supplementary Table 4 contains correlations of predicted motif pioneering strength to various enhancer positions. Supplementary Table 5 contains parameters used in equilibrium kinetics modeling. Supplementary Tables 6-7 contains parameters used in non-equilibrium kinetics modeling.

## Data and code availability

Raw and supplementary processed data for sequencing experiments have been deposited at GEO under series accession number GSE306105. The software used to train and interpret BPNet and ChromBPNet stemmed from the BPReveal software release ^51^ (v.4.0.4) and is available (https://github.com/mmtrebuchet/bpreveal/tree/4.0.4). Trained models, supporting intermediate data and motif mappings are available on Zenodo at (10.5281/zenodo.17187407). Code and analysis to process sequencing data, train models and produce analysis is available on GitHub (https://github.com/zeitlingerlab/Weilert_mESC_accessibility_2025).

## Acknowledgements

We thank Robb Krumlauf, Barak Cohen, Yuning Zhang, Neşet Özel, Sharon Torigoe and Žiga Avsec for helpful comments and suggestions. We also thank the Stowers Technology Centers for support: Sequencing and Discovery Genomics (Anoja Perera, Michael Peterson, Amanda Lawlor and Rhonda Egidy), Cytometry (Jose Javier, KyeongMin Bae and Jeff Haug), Genome Engineering (Kyle Weaver and Victoria Hassebroek) and Tissue Culture (Yan Wang, Sonia Ghosh, Maria Katt, Shilpa Waduwawara and Chongbei Zhao).

## Author contributions

M.W. and J.Z. conceived the project with further support from R.M.-C. All data processing, deep learning modeling and analysis work was performed by M.W. CRISPR mutations were conceived by M.W. and J.Z., while CRISPR was performed by K.D. Sequencing experiments were generated by K.J.B, K.D., S.K and H.J. Kinetics models were conceived and designed by R.M.-C. Interpretation of kinetics simulations were done by R.M.-C., J.Z. and M.W. The manuscript was prepared by M.W., R.M.-C. and J.Z. with input from all authors.

## Methods

### Cell culture

R1 ESCs were cultured on 0.1% gelatin-coated plates without feeder cells in N2B27 medium (DMEM/F12 with 1:1 mix of GlutaMax/N2 and Neurobasal medium/B27, Invitrogen) supplemented with 2 mM L-glutamine (Stemcell Technologies), 1x 2-mercaptoethanol (Millipore), 1x NEAA (Stemcell Technologies), 3 µM CHIR99021 (Stemcell Technologies), 1 µM PD0325901 (Stemcell Technologies), 0.033% BSA solution (Invitrogen) and 107 U/mL LIF (Millipore).

### CRISPR/Cas9 motif mutations

For each of the five candidate mutations across two genomic loci described below in mouse R1 ESCs, guide RNA target sites were designed using the IDT target predictor tool by evaluating the predicted on-target efficiency score and off-target potential. Alt-R CRISPR-Cas9 crRNA was designed to contain ∽40 bases of homology from the targeted cut site. Equimolar amounts (stock of 100 μm) of Alt-R crRNA and tracrRNA-ATTO550 were mixed to form at gRNA at a final concentration of 50 μM. The mixture was heated at 95°C for 5 minutes and cooled at RT. The single-stranded donor oligonucleotides (ssODN) were also designed to contain ∽40 bases of homology from the targeted cut site using the IDT software tool. A ribonucleoprotein (RNP) complex was formed by combining 150 pmol of gRNA (crRNA+tracrRNA) and 125 pmol of Cas9 HiFi v3 protein (IDT) with hybridization for 20 minutes at RT. The RNP was combined with 100 pmol of ssODN donor and 100 pmol of electroporation enhancer v2 and delivered to 1.5e5 cells by Neon electroporation (1,400 V, 10 m, 3 pulses; Neon Transfection System, MPK5000, Life Technologies). Immediately after electroporation, cells were cultured in 0.5 μM Alt-R HDR enhancer V2 of 0.69 mM. After 24 hours, cells were washed with PBS before FACS sorting on S6 FACSymphony. Single cells were directly sorted into 96-well plates. Cells were screened for the expected mutations through paired-end sequencing on an Illumina MiSeq instrument (250 cycles). On-target indel frequency and expected mutations were analyzed using CRIS.py.^108^ Only clones with the intentional mutation and sequence alignments >90% were chosen for future experiments. Close selections for each candidate mutation/locus are described below. Upon clone selection, ATAC-seq and ChIP-nexus experiments were prepared alongside wildtype R1 ESC control samples.

The first candidate set of mutations were located at the intronic *Btbd11* enhancer assessed in previous work.^36^ We mutated every pairwise combination of the predicted *Sox2* motif on chr10:85,539,634-85,539,643 (mm10) and the predicted low-affinity *Oct4-Sox2* motif on chr10:85,539,666-85,539,681 (mm10). This resulted in three total CRISPR mutations: (1) *Sox2* mutated, (2) *Oct4-Sox2* mutated and (3) both motifs mutated. In these cases, the *Sox2* motif was mutated from CCTTTGTTCC (wildtype) to CCTAGGTTCC (mutant) and the *Oct4-Sox2* motif was mutated from AATTATAATGATAAT (wildtype) to AATCATAAGGATAAT (mutant). One clonal cell line for each mutation was used to perform two biological ATAC-seq replicate experiments.

The second set of mutations were introduced at the upstream *Akr1cl* enhancer. We mutated one of the two *Sox2* motifs, located at chr1:65037654-65037663 (mm10) and chr1:65037753-65037762 (mm10). These two *Sox2* motifs contained the exact same motif sequence CCCTTTGTC (wildtype) and each was mutated into the motif sequence CCCTAGGTC (mutant), which differs by two bases. One clonal cell line for each mutation was used to perform two biological ATAC-seq replicate experiments.

### ATAC-seq experiments

For each ATAC-seq experiment, 100,000 R1 wildtype or CRISPR-edited mouse embryonic stem cells were used. Cells were washed once in cold PBS, with centrifugation at 500 x g and 4 °C for 5 min. The cell pellets were then resuspended in 50 µL of cold ATAC Resuspension Buffer (10 mM Tris-HCl pH 7.4, 10 mM NaCl, 3 mM MgCl2) supplemented with 0.1% IGEPAL CA-630, 0.1% Tween-20 and 0.01% Digitonin (Promega, G9441). After a 3 min incubation on ice, 1 mL of cold ATAC Resuspension Buffer supplemented with 0.1% Tween-20 was added and the samples were centrifuged at 500 x g and 4°C for 10 min. Tagmentation was performed at 1000 rpm and 37 °C for 30 min in a 50 μL reaction volume containing 10 μL of 5x Tagment DNA Buffer (50 mM Tris-HCl pH 7.4, 25 mM MgCl2, 50% DMF), 0.5 μL 10% Tween-20, 0.5 μL 1% Digitonin,1 µL of 20 µM Tn5 transposase loaded with oligonucleotides and 38 µL water. For all experiments, the Tn5 used was purified in-house using pETM11-Sumo3-Tn5 and His6-tagged SenP2 protease plasmids as performed previously.^21,24,36,109^ DNA fragments were purified using the Monarch PCR & DNA Cleanup Kit (NEB, #T1030) and used as input for library preparation using Illumina Nextera Dual Indexing. During library preparation, qPCR was performed to prevent over-amplification^109^ and libraries were purified using a 2% agarose gel and the Monarch DNA Gel Extraction kit (NEB, #T1020). Multiple, highly correlated biological replicates were performed for all ATAC-seq experiments and paired-end sequencing was performed on an Illumina NextSeq 500 and 2000 instruments (2 x 75 bp cycles).

### ChIP-nexus experiments

For each ChIP-nexus experiment, 10 million R1 wildtype or CRISPR-edited mouse embryonic stem cells were used, as described previously.^35,36^ In brief, the cells were first washed once with PBS, crosslinked using 1% formaldehyde and quenched after 10 min using 125 mM glycine. Fixed cells were then washed twice more with ice cold PBS and resuspended in chilled lysis buffer (15 mM HEPES pH 7.5, 140 mM NaCl, 1 mM EDTA, 0.5 mM EGTA, 1% Triton X-100, 0.5% *N*-lauroylsarcosine, 0.1% sodium deoxycholate and 0.1% SDS). After a 10 min incubation on ice, a Bioruptor Pico (Diagenode) was used to sonicate the chromatin with five or six cycles, 30 sec on/off. The commercially available Sox2 antibody (Active Motif, 39843) was used for these experiments. For each ChIP, 10 µg of antibody was conjugated to 50 µL of Protein A or G dynabeads (Invitrogen). Following the ChIP, the standard ChIP-nexus library preparation, including end repair, dA-tailing, adapter ligation, barcode extension and lambda exonuclease digestion, was performed as described,^110^ except for the following two changes: the ChIP-nexus adapter mix contained four fixed barcodes, and PCR library amplification was performed directly after circularization of the purified DNA fragments (without addition of the oligo and BamHI digestion). Single-end sequencing was performed on both Illumina NextSeq 500 and Illumina NextSeq 2000 instruments (75 or 150 cycles) at the Stowers Institute. For all experiments, at least two biological replicates were performed. The current ChIP-nexus protocol can be found on the Zeitlinger lab website at https://research.stowers.org/zeitlingerlab/protocols.html.

### ATAC-seq data processing

Paired-end ATAC-seq reads from a doxycycline-inducible knockout cell line ZHBTc4 stored under GSE174774 were reprocessed alongside the paired-end sequencing reads from ATAC-seq experiments conducted in this work. Reads were pre-trimmed for adapters using Cutadapt (v.4.2)^111^ and aligned to the mouse genome assembly mm10 using bowtie2 (v.2.3.5.1).^112^ Any CRISPR ATAC-seq experiments were aligned to a custom mm10 genome assembly modified to contain the expected CRISPR mutations. Duplicated alignments were marked using Picard (v.2.23.8).^113^ Alignments were then deduplicated, filtered for fragment lengths exceeding 600 bp, corrected for dovetailed alignment pairs and end-adjusted to accommodate the Tn5 enzymatic cut correction of -4/+4. Normalized read-per-million (RPM) ATAC-seq tracks were generated by scaling the aligned coverage to the weighted total reads as described previously.^21^ Cut site ATAC-seq coverage used for training ChromBPNet models were generated by isolating the bases on each end of an aligned fragment, consolidating those “cut sites” into coverage tracks. ATAC-seq peaks were mapped using MACS2 (v.2.2.7.1)^114^ with default settings. High-confidence peaks were then kept from the most reproducible replicate pair using the Irreproducible Discovery Rate framework (IDR) (v.2.0.3)^115^ for downstream analysis and model training. Whenever visualizing observed ATAC-seq cut site coverage at individual loci, we used Locally Estimated Scatterplot Smoothing (span=0.2)^116,117^ rather than ATAC-seq fragment pileup.

### ChIP-nexus data processing

Single-end sequencing reads from GSE137193 were reprocessed alongside the single-end sequencing reads from ChIP-nexus experiments conducted in this work. Reads were preprocessed by trimming off fixed and random barcode components from the sequenced reads, reassigning barcode information as read names for downstream processing. Next, reads were pre-trimmed for adapters using Cutadapt (v.4.2)^111^ and aligned to the mouse genome assembly mm10 using bowtie2 (v.2.3.5.1).^112^ Any CRISPR ChIP-nexus experiments were aligned to a custom mm10 genome assembly modified to contain the expected CRISPR mutations. Alignments were then deduplicated from aligned coordinates and their corresponding barcodes. Coverage was generated pooling the first “stop” base of each deduplicated read, separated into two tracks by read orientation. Normalized read-per-million (RPM) ChIP-nexus tracks were generated by scaling the aligned coverage to the weighted total reads as described previously.^21^ ChIP-nexus peaks were mapped using MACS2 (v.2.2.7.1)^114^ with parameters designed to resimulate the full fragment length coverage rather than the single stop base coverage (--keep-dup=all -f=BAM --shift=-75 --extsize=150). High-confidence peaks were then kept from the most reproducible replicate pair using the Irreproducible Discovery Rate framework (IDR) (v.2.0.3).^115^ Peaks were then filtered for non-standard chromosomes, sequence ambiguities and presence on chromosome boundaries. After filtering, 154,827 peaks were used for downstream analysis and model training.

### TT-seq data processing

Paired-end TT-seq reads from a doxycycline-inducible knockout cell line ZHBTc4 stored under GSE174774 were reprocessed. Using STAR (v.2.7.3a)^118^ on default settings, transcript reads were both aligned to the mouse genome assembly mm10 and collected into gene counts tables derived from Ensembl v.98 gene annotations.^119^ Alignments were then deduplicated, filtered for fragment lengths exceeding 600 bp and corrected for dovetailed alignment pairs. We generated strand-separated genome-wide coverage using the by isolating the 5’ ends of each aligned read, thus representing each single-base read as an “event of transcription”, rather than fragment pileup. Normalized read-per-million (RPM) TT-seq single-base tracks were generated by scaling the stranded, aligned coverage to the weighted total reads as described above for ATAC-seq and as done previously.^21^

### Model training and validation

BPNet is a convolutional network designed to learn genomic data from input DNA sequences alone.^36^ Using the BPReveal implementation^51^ (v.4.0.4) of BPNet’s previously described architecture, we trained a multi-task model to learn the ChIP-nexus binding of Oct4, Sox2, Nanog, Klf4 and Zic3 signals. Input DNA sequences were derived from the sequences of 154,827 IDR-reproducible peak regions, described above. Models used for downstream analysis were validated on 27,866 sequences from chromosomes 2/3/9, tested on 25,351 sequences from chromosomes 1/8/17 and trained on 101,451 sequences from the remaining standard chromosomes. Post-optimization, the final BPNet binding model was trained with 8 convolutional layers, a filter depth of 128, an input filter length of 7 and an output filter length of 7. Model performance was determined based on Spearman correlations of predicted versus observed total experimental counts, Jensen-Shannon distances of predicted and observed profiles and multinomial negative log-likelihoods of predicted and observed profiles. With this associated architecture, input DNA sequences were 2032 bp long, resulting in the desired output prediction window of 1000 bp, as BPReveal’s reimplementation of BPNet does not pad through convolutions, but rather requires a larger input sequence to provide the receptive field with sufficient sequence information as the model deepens. Model stability was cross-validated by training two other models with different training, validation and test regions and assessed via described performance metrics alongside consistently learned sequence patterns upon application of interpretation tools DeepSHAP^52,53^ and TF-MoDISco.^36,54^

ChromBPNet is a modified training approach to BPNet, applying BPNet convolutional architecture to learning ATAC-seq data in a two-step model training process to eliminate the positional Tn5 sequence biases of ATAC-seq experiments.^38^ Using the BPReveal implementation^51^ (v.4.0.4) of ChromBPNet, we trained the first “Tn5 bias” model on 22,053 inaccessible, low-count reads from our pooled ATAC-seq experimental coverage in wildtype mESCs, using the same training, validation and test chromosomes described above for the final BPNet binding model. We confirmed that this bias model did not learn TF binding motifs and that only the expected Tn5 bias sequence patterns were returned through assessing model predictions upon *in silico* motif injection alongside application of interpretation tools DeepSHAP^52,53^ and TF-MoDISco.^36,54^ We next trained a second, residual model alongside the now-frozen bias model to explain bias-removed ATAC-seq accessibility using the same 154,827 sequences split between the same training, validation and test sets used in the BPNet model training, described above. These regions were confirmed to contain an appropriately diverse range of accessibility to train a robust ChromBPNet model. During this training step, we considered only the positional information provided by the bias model when assessing output predictions. We optimized ChromBPNet model architecture using the same performance metrics described above for the BPNet binding model and post-optimization, the final residual ChromBPNet model was trained with 8 convolutional layers, a filter depth of 128, an input filter length of 7 and an output filter length of 7. With this associated architecture, input DNA sequences were 2032 bp long, resulting in the desired output prediction window of 1000 bp. Model stability was assessed with cross-validation training, performed identically to BPNet models described above.

To capture accessibility rules after depletion of Oct4, a separate ChromBPNet model was trained for each of the six time points (0h, 3h, 6h, 9h, 12h and 15h) from the reprocessed GSE174774 ZHBTc4 ATAC-seq data in mESCs, using the same bias model, optimized architecture and cross-validated region sequence sets as the original mESC R1 ChromBPNet model. Model performance and stability were determined using the same performance metrics described above for the BPNet binding model and the same cross-validation training approach. After confirming that the models were stable and performed well, we also validated that the sequence rules and motifs returned by the initial timepoint (0h) ChromBPNet model of the ZHBTc4 cell line were comparable to our wildtype R1 ChromBPNet model.

### Motif mapping

For all trained ChromBPNet residual accessibility and BPNet binding models, we generated sequence contribution scores using BPReveal’s implementation^51^ (v4.0.4) of DeepSHAP, modified to generate hypothetical contribution scores akin to those from DeepLIFT.^52,53^ Using counts contribution scores (explaining the total amount of binding or accessibility across a region) inputted to the tfmodisco-lite implementation of TF-MoDISco^36,54^ (v2.2.0, https://github.com/jmschrei/tfmodisco-lite), we generated contributing motif representations called CWMs (contribution weight matrices) for each trained model. We manually curated motif/TF identities based on existing literature, returning expected pioneer motifs and motifs important for TF binding in mESCs and validating the cross-validated models returned the same motif representations. It should be noted that while the *Oct4* motif was discovered, this motif was not robustly discovered across model folds for either binding or accessibility, produced poor Oct4 ChIP-nexus footprinting across mapped motifs and was often enriched in ERV regions. Thus, we excluded the *Oct4* motif from further downstream analysis.

To map the TF-MoDISco CWM motif representations back to the genome, we performed contribution weight matrix scanning (CWM scanning) implemented by BPReveal^51^ (v4.0.4), as described previously.^36^ Briefly, across all 154,827 of regions considered above, we mapped a motif if the genomic region matched criteria designated by the corresponding TF-MoDISco motif distributions. The criteria used in motif mapping here required that (1) the Jaccardian similarity between the motif CWM and the genomic site’s sequence contribution exceeded the 20th percentile of the TF-MoDISco motif representations and (2) the contribution L1 magnitude is higher than the TF-MoDISco motif seqlets lowest contribution score. Unlike the previous approach^36^, we did not require that the log-odds score relative to the TF-MoDISco motif PWM be greater than zero. This allowed us to rely solely on contribution similarity and magnitude, to encourage our mapping strategy to allow very degenerate sequences, provided they possess sufficiently well-positioned counts contributions across the motif coordinate bases. Upon CWM scanning, we refined the mappings by resolving redundant (e.g. *Oct4-Sox2* and *Sox2*) and palindromic mappings. For each motif, its percentile relative to the original seqlet distributions of CWMs (Jaccardian similarity) and PWMs (information content) was calculated. We note here that unless otherwise explicitly stated, PWM scores refer to the information content score and not the log-odds score.

We performed PWM-scanning in order to benchmark our contribution-derived mapping approach using the FIMO tool from MEMESuite^59,60^ (--skip-matched-sequence --max-strand --thresh 0.001) with TF-MoDISco defined seqlets as the consensus frequencies across only our 154,827 regions of interest. Thresholds of significant matches (*p*=0.001) were set low enough to return common and poor mappings.

### Isolation and context scores

Previous work has shown that predicting the log-fold-change effects of an injected TF binding motif over a random *in silico* background using BPNet can result in a strong proxy for motif affinity.^68^ Referred to here as the “isolation” scores of an injected sequence, this injection strategy removes the surrounding genomic context of a mapped motif to measure its intrinsic effects out of context. Briefly, a sequence was injected into randomly generated 256 background sequences, predictions were measured with and without the injected sequence, then the log-fold-change was calculated between averaged injected and background levels. Background sequences were derived by sampling trained peak regions to match the baseline CpG enrichment and GC-content of the mouse genome, then di-nucleotide shuffling the underlying sequences. CpG ratios were calculated as (GC-content / 2)^2^ and normalized to each region width, as previously described.^120^

In contrast to isolation scores, the “context” scores of a mapped motif involves measuring a motif’s effect within its genomic context rather than in isolation, as described previously.^21^ Mapped motif sequences were mutated via replacement with random insertions. Across 16 trials of mutations, average mutation response was calculated, then log-fold-change was computed between mutated and wildtype profiles.

### Cooperativity characterization

We quantified a pair of mapped motifs as cooperative when they produced accessibility greater than the sum of their individual parts. More precisely, for each pair of mapped motifs we predicted the accessibility effects across the *in vivo* wildtype sequence (AB), sequences with one motif mutated (A or B) and sequences with both motifs mutated (null). We characterized accessibility effects as the total predicted ATAC-seq reads within the surrounding 1 kbp region. These total predicted reads were exponentiated from the log-space counts outputs of BPReveal models. As previously described^19^, “joint” effects of paired motifs were characterized as (AB - null). “Marginal effects” of individual motifs were characterized as ((A - null) + (B - null)). Cooperativity scores were assigned by taking the log-fold-change of the joint effects over the marginal effects for all mapped motif pairs. When testing whether a specific type of motif pair was significantly cooperative, we used a one-tailed Wilcoxson test comparing whether mapped motif pairs belonging to that type arranged at specified distances (typically bins of 10 bp) had greater cooperativity scores than mapped motif pairs at distances that were deemed too long to correspond to cooperative pioneering (greater than 400 bp), with multiple-comparison Bonferroni corrections and an adjusted significance cutoff requirement of *p* < 10^-7^.

### PBM comparisons

Processed PBM experiments that were close matches to our TFs of interest were downloaded from UniPROBE^64^ in order to compare the *in vitro* binding Z-scores of each measured 8-mer sequence to the PWM and isolation scores of our TF/motif pairs. To compare PBM Z-scores to either mapped motif PWM scores or isolation scores, we trimmed the mapped motif sequences down to the expected “core” 8-mer sequences. If this trimming resulted in multiple redundant 8-mers, we took the one with the maximum isolation score (PWM scores would be identical). When 8-mer sequences were redundant within the PBM datasets due to reverse complement matches, we took the maximum Z-score. We compared human Sox2 PBM data to our Sox2/*Sox2* TF/motif pair.^61^ We compared mouse Klf1 PBM data to our Klf4/*Klf4* TF/motif pair.^62^ We compared human Zic3 PBM data to our Zic3/*Zic3* TF/motif pair.^63^ We compared human POU5F1 PBM data to our Oct4/*Oct4-Sox2* TF/motif pair, favoring the *Oct4* motif component of the heterodimer *Oct4-Sox2* motif when trimming down 8-mers.^61^

### Differential enrichment

In order to measure differential enrichment of accessibility and enhancer RNA across changing Oct4 concentrations, we performed DESeq2^90^ on the reprocessed ATAC-seq and TT-seq data derived from a time course experiment with decreasing Oct4 levels after doxycycline withdrawal.^67^ DESeq2 was performed in all cases with default parameters and FDR=0.05. For each DESeq2 model described below, we conducted pairwise comparisons between the wildtype time point (0h) and all corresponding mutant time points (3h-15h), computing the log2(mutant/wt) values for each. As we previously described^21^, log2(mutant/wt) < 0 will represent a loss in signal enrichment in the mutant, while log2(mutant/wt) > 0 represents a gain in enrichment in the mutant.

More specifically, to measure differential accessibility across accessible regions of interest over changing Oct4 concentrations, we built a DESeq2 model containing ATAC-seq cut site coverage for every replicate and time point within the 154,827 genomic regions (see “Model training and validation”) used for model training. To measure differential enhancer expression over changing Oct4 concentrations, we used the same procedure described above for differential accessibility. We collected base-resolution TT-seq coverage across the 154,827 accessible regions of interest for every replicate and time point, building a DESeq2 model from the collected coverage.

### Nucleosome-mediated TF modeling

Nucleosome-mediated TF cooperativity has often been modeled under thermodynamic equilibrium,^27,77,79^ where the chromatin exists in two conformations, closed (with nucleosome) and open (without), with basal transitions between the closed and the open conformations. In these models, a TF promotes the open conformation when the affinity of the TF is higher in the open conformation than in the closed conformation. Given the assumption of thermodynamic equilibrium, this implies an equivalent reduction in the affinity of the nucleosome in the open chromatin conformation relative to the closed conformation, thereby accounting for a competition between the TF and the nucleosome. This model contradicts however the concept of a pioneer TF, whose ability to open chromatin depends on its ability to bind the nucleosome in the closed state.^18^ Intuitively, pioneering TFs should produce strong opening effects that do not entirely rely on higher affinity binding in the open state. To achieve this, we chose to consider a kinetic model, a two-conformation TF-nucleosome model away from equilibrium. This allowed us to separate the effect of the nucleosome on TF binding, which can be small, from the opening effect of the TF, which can be strong.

For the single motif case, we consider a model with 4 states: (1: closed, unbound), (2: closed, bound), (3: open, unbound), (4: open, bound), with reversible transitions between states 1-2, 1-3, 2-4, 3-4 (Figure S5B). In order to account for 2 TF motifs, the model is extended to 8 states, accounting for all possibilities of TF and nucleosome bound or not, and the corresponding transitions (Figure 5C). In both cases, the system is assumed to evolve over time according to Markovian dynamics, which settles into a steady-state. To quantify accessibility, we consider the sum of the steady-state probabilities of the open states, here represented as states 3 and 4 for the single TF model (Figure S5B). Such steady-state probabilities can be calculated by solving for the null space of the transition rate matrix (formally, the Laplacian of the graph associated with the system). For a model with N states, this is a square matrix of size N x N where the value of the entry in position i,j contains the transition rate from state j to i if i ≠ j, and the diagonals contain the negative of the sum of the rest of the column.^121,122^ For a given set of parameters, we obtain a basis for the nullspace of this matrix using SciPy (v.1.12.0, scipy.linalg.null_space)^123^ and obtain the steady-state probabilities by normalizing each entry by the total sum, such that the probabilities sum to one.

As we saw, a TF has motifs of different affinities, whose effective affinity *in vivo* can be modulated by the surrounding sequence context. Similarly, the effect of the TF once bound at the motif may also depend on surrounding molecules. Therefore, in order to explore how the model behaved for a single TF motif and for two TF motifs, we sampled parameter sets accounting for potential variation in the motif binding affinity and effect of the TF(s) once bound.

For a given parameter set, we consider that the TF binding rate (k_on_) is independent from whether or not the motif or the nucleosome is present. This is in line with an assumption for a diffusion-limited binding rate. TF concentration is absorbed in the TF association rate (k_on_, such that k_on_=k’_on_ [A], with [A] the concentration of TF A, such that changing concentrations is simulated by changing k_on_). Similarly, we consider motif-independent basal opening rate (k_open_) and closing rate (k_close_), and the effect of the TF on these rates, which are modulated by a factor o for the opening rate and c for the closing rate when the TF is bound. We select a value for the unbinding rate of the TF from the open conformation in the absence of TF (non-specific unbinding rate on naked DNA, k_off_). This unbinding rate is assumed to be reduced by a factor g in the presence of a motif. The destabilizing effect of the nucleosome on the TF is also introduced in the unbinding rate, which is increased by a factor (D_n_) in the absence of motif, and a factor (D_m_) in the presence of a motif. Given these parameters, we can calculate the steady-state probability of the accessible conformation for the parameters corresponding to the motif being present, and for those corresponding to the motif being absent, such that we can calculate the log-fold-change in accessibility. This provides an equivalent metric to the isolation scores derived from deep learning predictions of sequence injections *in silico*.

For two TFs A and B, a similar approach applies where we expand the model to consider four sequence scenarios: no motifs, motifs A and B, motif A only and motif B only. When two TFs are bound, we must make an assumption for what o_AB_ and c_AB_ are. If o_A_ is the factor increase in the opening rate when TF is bound at motif A and o_B_ is the factor increase in the opening rate when TF is bound at motif B, we choose o_AB_ = o_A_ x o_B_ for the opening rate when both are bound. The same multiplicativity assumption is made for the joint effects on the closing rate c_AB_. This multiplicativity arises naturally if we assume that there is an activation energy E associated to a transition rate k, with each TF having an additive effect on this activation energy x_A_ or x_B_. Following the Arrhenius equation:

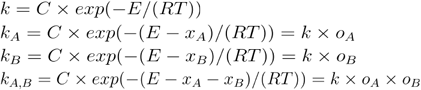

When simulating pioneering TF binding to its respective motif, we typically chose to model Sox2 binding to the *Sox2* and *Oct4-Sox2* motifs. We chose parameters from plausible biological ranges, described below, depicted graphically (Figure S5B) and reported each parameter regime for every simulation conducted in Supplemental Tables 5-7.

For TF binding on rates (k_on_), we assume a diffusion-limited binding on rate. This rate is therefore unaffected by whether or not there is a nucleosome and unaffected by whether or not there is a motif. Diffusion-limited binding rates for TFs have been estimated to be on the order of 10^-2^ - 10 s^-1^nM^-1^.^124^ For Sox2, an *in vitro* binding constant for the DNA binding domain has been measured to be 0.75 x 10^-3^ s^-1^nM^-1^.^88^ The Sox2 concentration in mESCs is on the order of 60 - 120 nM,^85^ which would correspond to an apparent binding rate of 0.045 - 0.09 s^-1^. In line with these numbers, the apparent k_on_ for Sox2 has been previously estimated using single molecule tracking^87^ to have a value of 0.27 s^-1^. Therefore, we typically chose an intermediate of k_on_ = 0.1 s^-1^, which we kept fixed throughout all simulations.

Given that k_on_ is fixed for a given concentration, it is the TF binding off rates (k_off_) that account for differences in affinity due to nucleosome or motif presence. For TF binding off rate from the open conformation without motif (k_off,4≠3,no motif_), indicating a non-specific off rate, we chose a value of k_off,4≠3,no motif_ = 1 s^-1^, in line with previous observations.^87^ For TF binding off rate from the open conformation with the motif ((k_off,4≠3,motif_), this off rate is reduced by a factor g taken to be between g = [0.001, 0.09]. These values are aimed to cover very low-affinity motifs (high g ∼ 1), stronger motifs which have been measured *in vitro* to have Sox2 binding affinities of up to ten fold that of naked DNA (g = 0.1),^86^ and potentially higher motif affinities that could arise through binding cooperativity with TFs that we do not model explicitly (lowest g ∼ 1). For TF binding off rate from the nucleosomal conformation without motif (k_off,2≠1,no motif_), this off rate is increased by five fold relative to that for the open conformation,^86^ specified by □_no motif_ (or □_n_) in the graphical depiction (Figure S5B). For the TF binding off rate from the nucleosomal conformation with motif (k_off,2≠1,motif_), specified by □_motif_, this off rate is decreased by a factor taken from the range □_motif_ = [2, 10].

We assumed a basal chromatin opening rate (k_open_), in the absence of a bound TF and independently of whether there is or not the motif, to be k_open_ = 0.05 s^-1^, in the order of tens of seconds. To choose the basal closing rate(k_close_), we defined the nucleosome dissociation constant *d*, which we chose from a given range, and set k_close_ = k_open_ / d. Note that in the absence of a bound TF, the probability of the system being open is d / (1 + d) ∼ d for d << 1. We typically chose a range for d = [0.001, 0.05) to model a predominantly closed region in the absence of a bound TF. With the TF bound, the chromatin opening rate (k_open_) is increased by a factor o, and the chromatin closing rate is decreased by a factor c, which are reported in Supplemental Tables 6-7.

## Supplemental figures

**Supplemental Figure 1.**
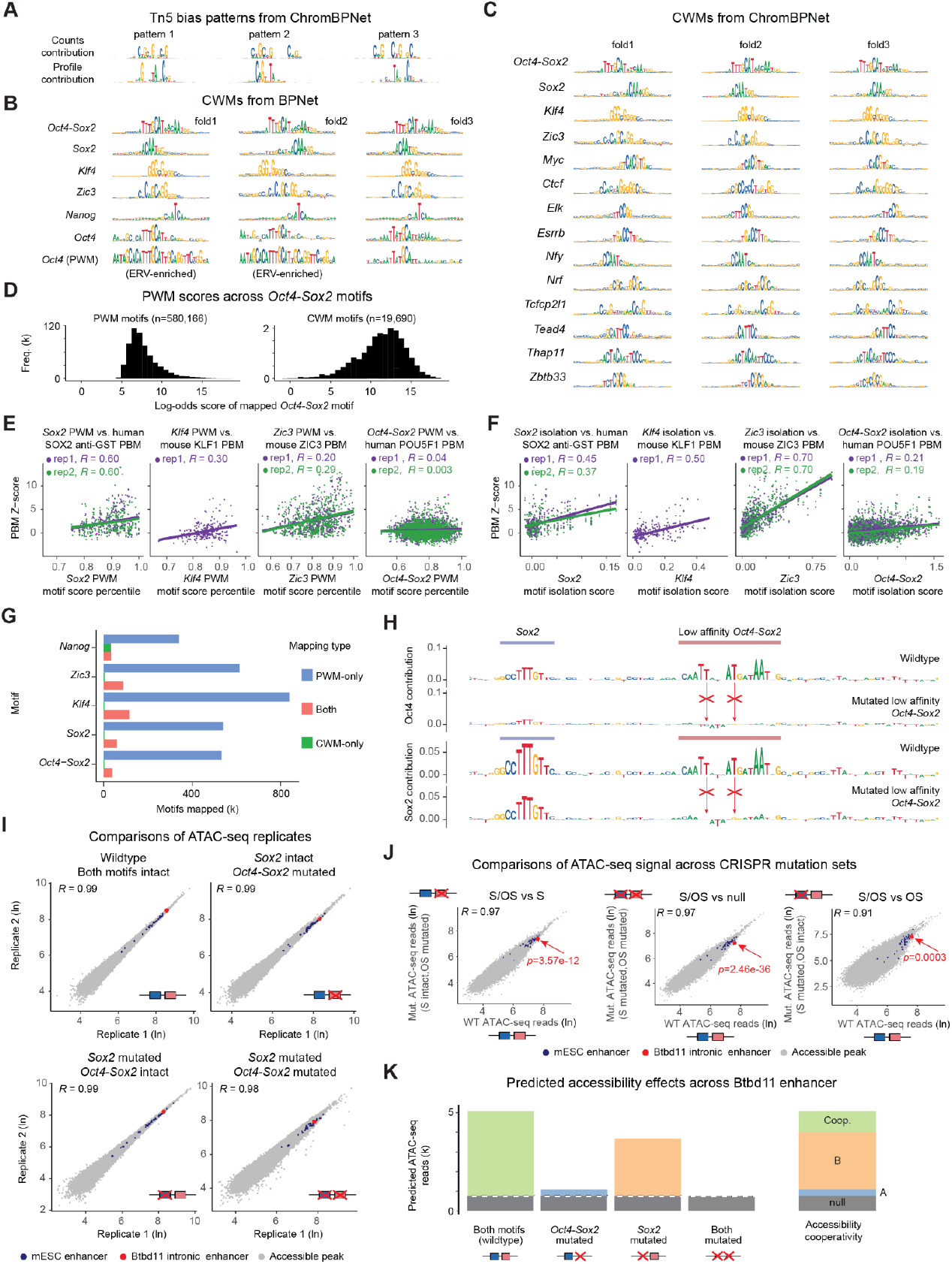
**A)** The ChromBPNet Tn5 bias model learned expected bias motifs, shown as PWMs for the top three patterns, summarized by TF-MoDISco from sequences with high counts contribution or profile contribution scores. **B-C)** Motifs shown as CWMs learned by **(B)** the BPNet binding model and **(C)** the ChromBPNet accessibility model across different folds as summarized by TF-MoDISco. While most motifs were robustly discovered in all folds, the single *Oct4* motif was not robustly discovered, often returning ERV-enriched mappings as these are recognizable in the *Oct4* PWMs (bottom row). **D)** Log-odds scores of *Oct4-Sox2* motifs learned by the accessibility model (right, n=19,960) show a broader distribution of affinity scores compared to *Oct4-Sox2* motifs mapped using PWM scanning (left, n=580,166). **E-F)** Scatterplots showing a relationship between model-derived motif features and available *in vitro* measured PBM binding data for human SOX2, mouse KLF1, mouse ZIC3 and human POU5F1 (OCT4), the TFs in the UniPROBE’s database that best match our TFs of interest. The PBM binding data are summarized as Z-scores for the 8-mer sequences that match the *Sox2, Klf4, Zic3* and *Oct4-Sox2* motifs. **(E)** The PWM score percentiles of all uniquely mapped motif sequences and **(F)** the corresponding binding isolation scores both correlate with the PBM Z-scores, as shown by Spearman correlation scores. PBM replicate experiments are shown in separate colors. **G)** Comparison of the number of motifs discovered only by PWM scanning, only by CWM scanning or both, using a low-stringency cutoff for PWM scanning necessary for discovering low-affinity motifs. As expected, PWM scanning discovered vastly more motifs. Motifs discovered by CWM scanning were typically a subset of the motifs discovered by PWM scanning, except for *Nanog* motifs, which are more difficult to map with PWM scanning. **H)** The intronic *Btbd11* enhancer with contribution scores from the BPNet binding model for Oct4 and Sox2 across the wildtype sequence (top) and mutated sequence (bottom). This shows that mutating the low-affinity *Oct4-Sox2* motif abolishes the predicted binding of Oct4 and Sox2. Note that the *Sox2* motif on the left is still predicted to be bound, although with slightly lower contribution in the mutant. **I)** ATAC-seq replicate comparisons show that the ATAC-seq experiments on wildtype and CRISPR clones mutating the *Btbd11* enhancer are reproducible (Spearman correlation coefficients reported on the scatterplots). Total reads were measured across ATAC-seq peaks found in wildtype experiments. **J)** Direct comparisons of ATAC-seq reads between wildtype and CRISPR *Btbd11* mutant experiments. As expected, the Spearman correlation coefficients are overall high but show differential accessibility across the *Btbd11* region, as calculated by DESeq2 with adjusted p values (red arrows). **K)** Accessibility across the *Btbd11* region as predicted by the ChromBPNet model for wildtype and after the *Sox2* and/or low-affinity *Oct4-Sox2* motif sequences were mutated. Changes were measured across coordinates chr10:85539400-85539800. To the right, predicted cooperative effects (green) are stacked upon individual marginal effects of motifs.

**Supplemental Figure 2.**
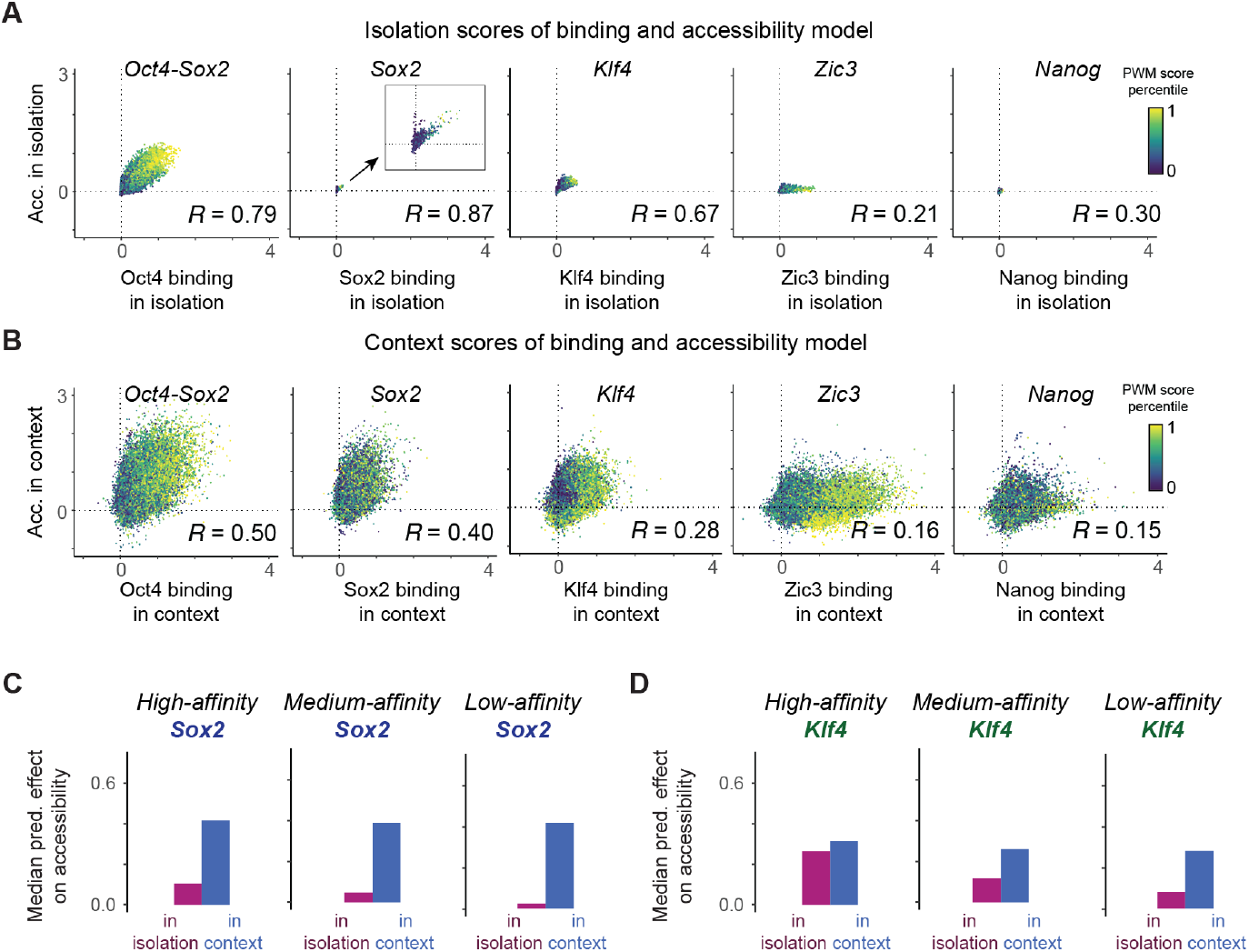
**A-B)** Comparison of the **(A)** isolation scores or **(B)** context scores obtained from the binding model (x-axis) and the accessibility model (y-axis) for the *Oct4-Sox2, Sox2, Klf4, Zic3* and *Nanog* mapped motifs, colored by their PWM score percentiles. **C-D)** For all sets of motifs motifs mapped by only accessibility models, median isolation and context scores predicted by the accessibility model for **(C)** *Sox2* motifs and **(D)** *Klf4* motifs of high, medium and low affinity motifs (5k each, based on PWM score) show that low-affinity motifs receive a bigger boost in context than high-affinity motifs.

**Supplemental Figure 3.**
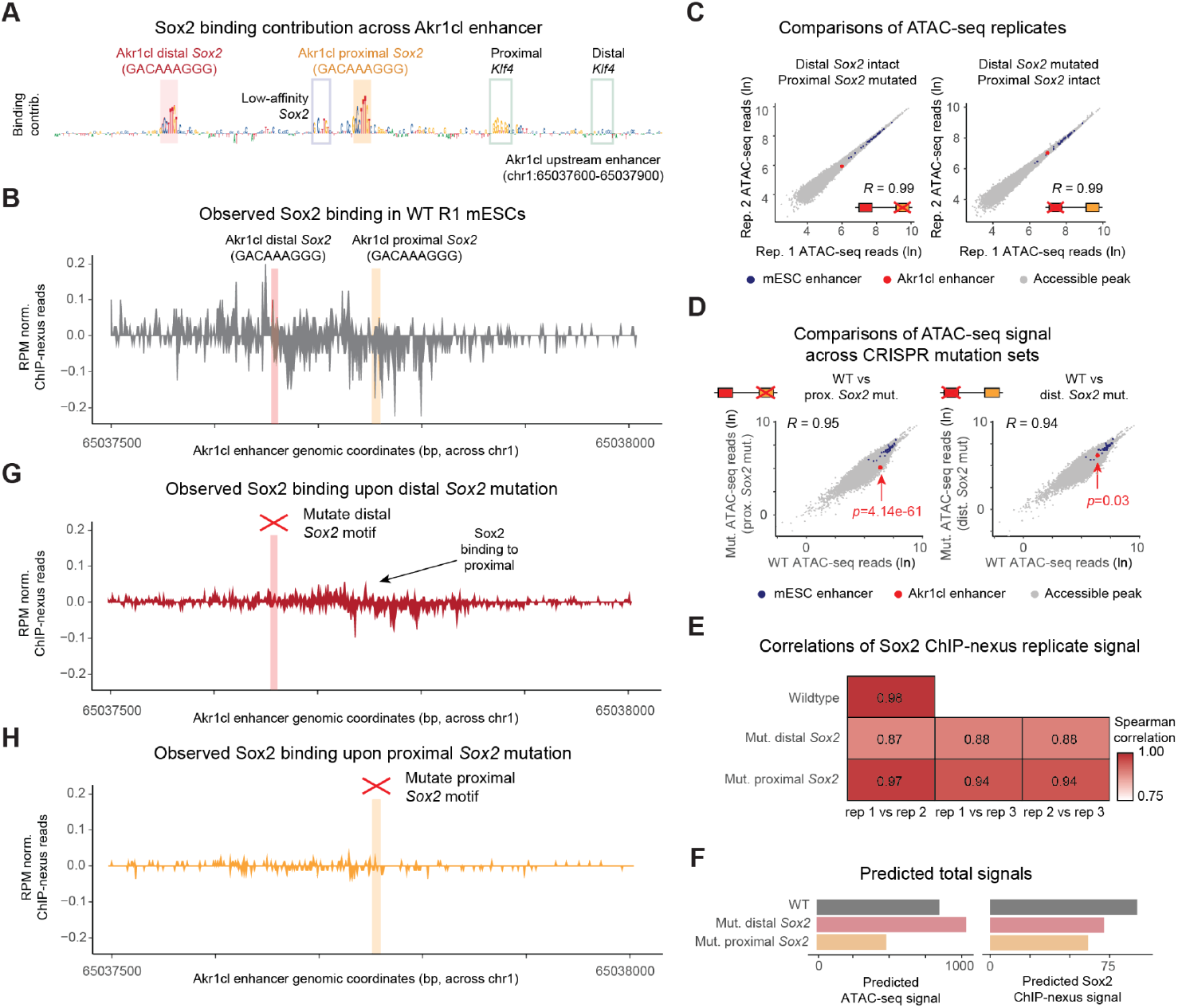
**A)** Sox2 binding contribution scores of the Akr1cl upstream enhancer (chr1: 65037600-65037900) with contributing motifs marked. **B)** RPM-normalized Sox2 ChIP-nexus across the Akr1cl enhancer, denoting distinct Sox2 binding profiles at both *Sox2* motifs. **C)** Replicate comparisons measuring total ATAC-seq reads across reproducible ATAC-seq peaks from wildtype and CRISPR *Akr1cl* mutant experiments. Spearman correlation coefficients are reported on scatterplots. **D)** Pairwise comparisons of pooled ATAC-seq reads from wildtype and CRISPR Akr1cl mutant experiments. Spearman correlation coefficients are reported on scatterplots. Differential accessibility between wildtype and each CRISPR Akr1cl mutant experiment was calculated using DESeq2, with adjusted significance values reported for the Akr1cl enhancer (red arrows). **E)** Replicate Spearman correlations measuring total Sox2 ChIP-nexus reads across reproducible ATAC-seq peaks from CRISPR *Akr1cl* mutant experiments. **F)** Predicted ATAC-seq signal or predicted Sox2 ChIP-nexus signal occurring across chr1: 65037600-65037900 across wildtype (black), distal *Sox2* mutant (red) and proximal *Sox2* mutant (orange) sequences. **G-H)** RPM-normalized Sox2 ChIP-nexus profiles across the *Akr1cl* enhancer from **(G)** the distal *Sox2* mutation and the **(H)** the proximal *Sox2* mutation CRISPR experiments.

**Supplemental Figure 4.**
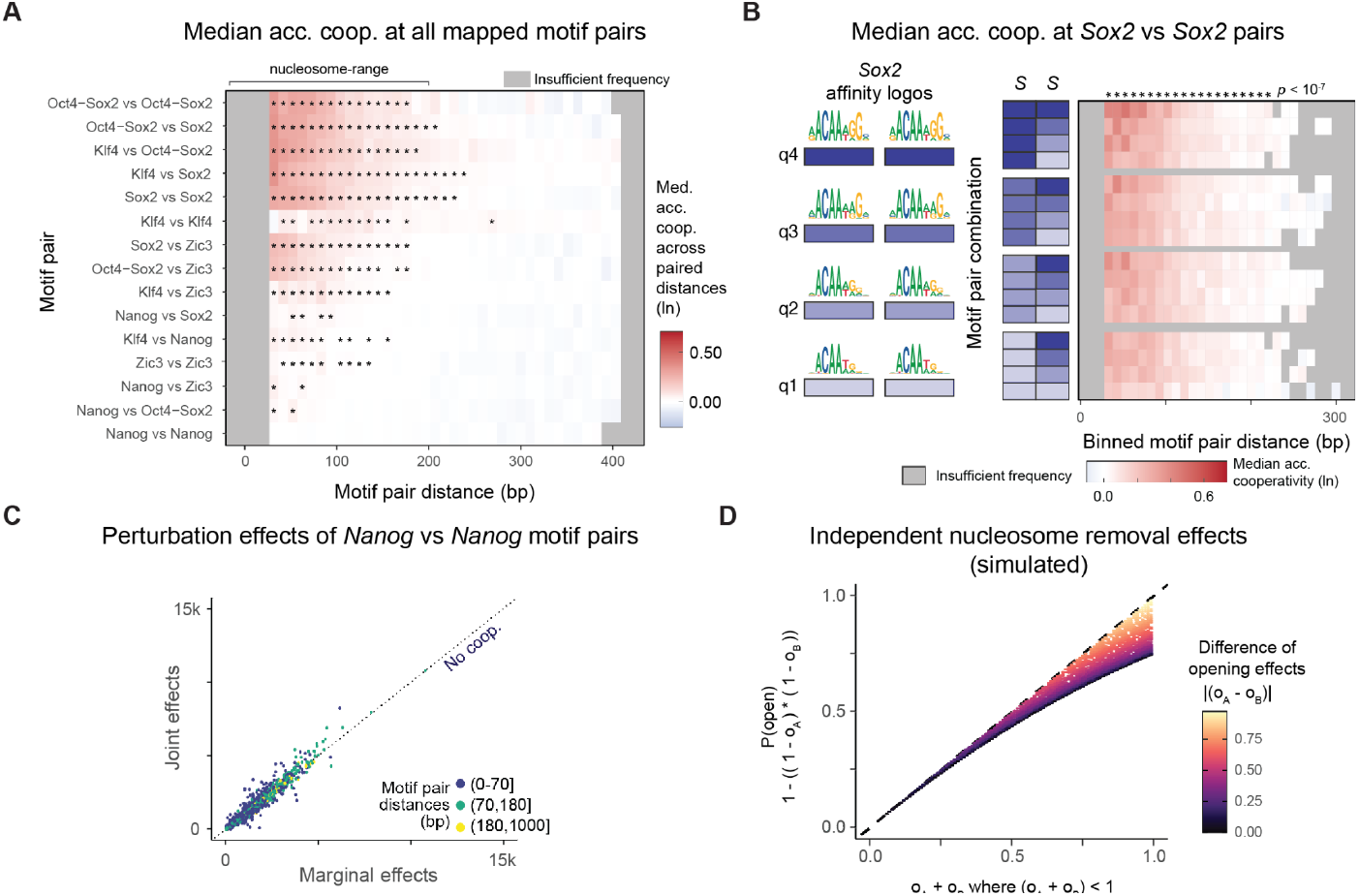
**A)** Median accessibility cooperativity across all motif pairs of key mESC motifs, separated by motif pair distances. Motif/distance combinations with fewer than 20 motif pairs were not considered (gray boxes). Cooperativity significance was derived from a one-tailed Wilcoxson test comparing whether motif pairs arranged at select binned distances were more cooperative than motif pairs arranged at very long distances (>400 bp) with multiple-comparison Bonferroni corrections and an adjusted significance cutoff of *p* < 10^-7^. **B)** Median accessibility cooperativity between *Sox2*/*Sox2* motif pairs across quartiles of motif affinity and motif pair distances, binned to 10 bp. Motif/distance combinations with fewer than 20 motif pairs were not considered (gray boxes). Cooperativity significance was derived from a one-tailed Wilcoxson test comparing whether *Sox2*/*Sox2* motif pairs arranged at select binned distances were more cooperative than *Sox2*/*Sox2* motif pairs arranged at very long distances (>400 bp) with multiple-comparison Bonferroni corrections and an adjusted significance cutoff of *p* < 10e^-7^. **C)** Comparison of joint and marginal predicted accessibility effects for the non-pioneer *Nanog*/*Nanog* motif pairs, colored by the pairs’ relative center-to-center distances to one another. **D)** Opening likelihoods for two pioneer motifs (o_A_ = [0.01, 1] and o_B_ = [0.01, 1] where o_A_ + o_B_ < 1) were independently sampled (n=10,000) and simulated based on the premise that each pioneer TF independently removes the same nucleosome with a certain probability (P(open) = 1 - ((1 - o_A_) * (1 - o_A_))), producing less than additive effects.

**Supplemental Figure 5.**
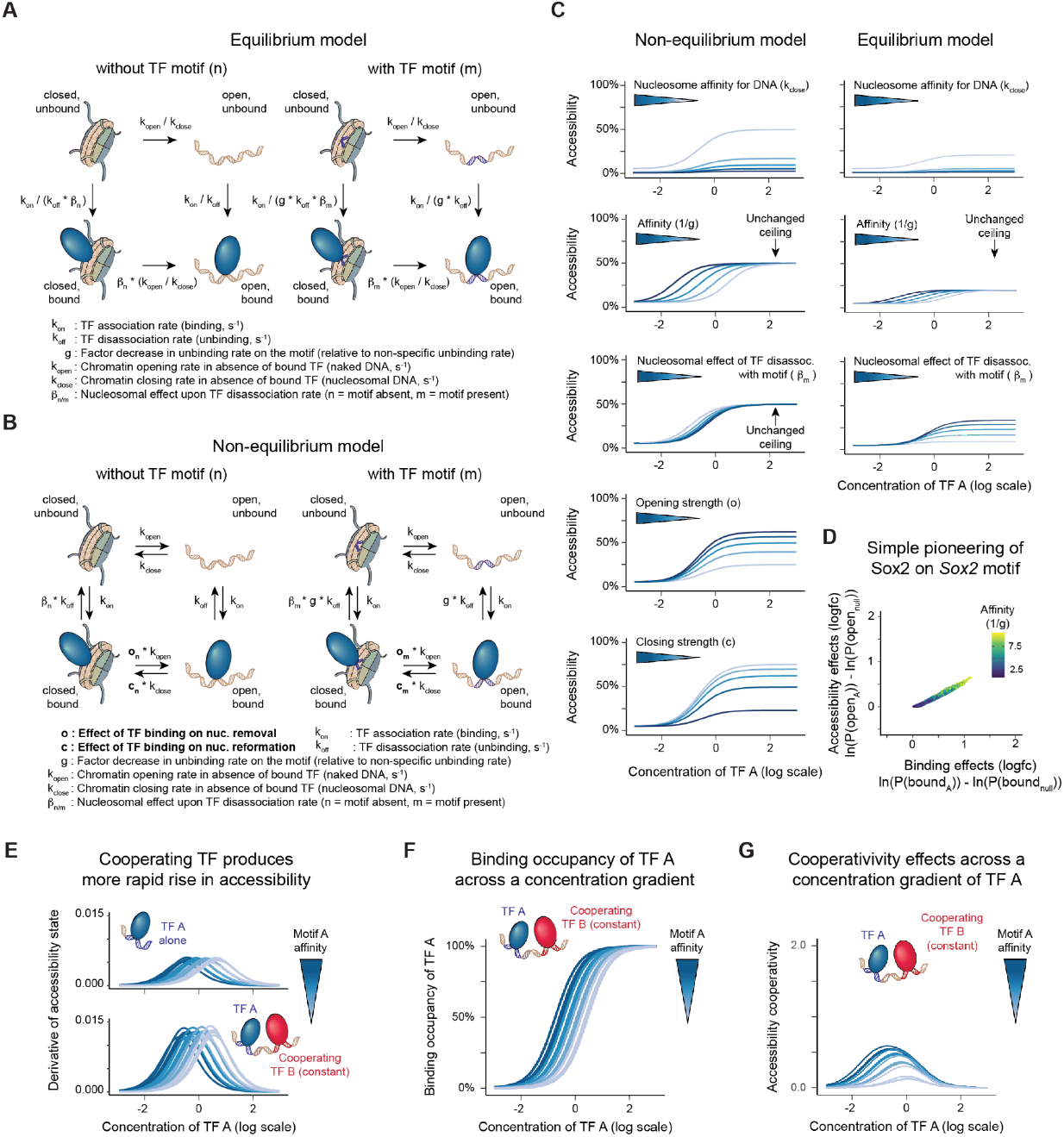
**A)** Parameterized graphic depicting a kinetic model in equilibrium of bound TF (called A) effects on nucleosome removal and reformation. Transitions are reversible, but they are only shown in one direction given that the equilibrium steady-state is only determined by the ratios between forward and backward rates, as indicated. On the left, the motif is not present and therefore recruits only non-specific binding to the TF. On the right, the motif is present and recruits specific TF binding, with motif affinity effectively given by the factor g. **B)** Parameterized graphic depicting a kinetic model out of equilibrium of bound TF effects on nucleosome removal and reformation. Parameter definitions are the same from graphical representation in **(A)**. Additional non-equilibrium parameters include o (effect of bound TF on nucleosome removal) and c (effect of bound TF on nucleosome reformation. Note that a higher value of c means less of an effect). **C)** 1 TF kinetic model simulations (n=1) in equilibrium (left) and non-equilibrium (right) showing the accessibility state over changing concentrations for a range of changing model parameters. We tested ranges of nucleosome intrinsic affinity for DNA, motif affinity, nucleosome effect upon TF disassociation in the presence of a motif, and in the case of the non-equilibrium model, TF binding effects on nucleosome removal and reformation. For the equilibrium model, the parameters were set to simulate moderate pioneering in the presence of a nucleosome with low intrinsic affinity to DNA. Parameter ranges can be found in Table S5 (equilibrium model) and Table S6 (non-equilibrium model). **D)** 1 TF kinetic model (out of equilibrium) simulations (n=50,000) showing the correlation between accessibility state, binding occupancy and motif affinity in the presence of a pioneer TF. Parameter ranges can be found in Table S6. Note that (effect of TF on nucleosome reformation), indicating that the simulated simple pioneering does not allow reduction of nucleosome reformation as a mechanism to increase accessibility, only nucleosome removal. **E)** Derivatives of accessibility curves from the 1 TF kinetic model (Figure 5B) and the 2 TF kinetic model (Figure 5E) showing that the cooperative (2 TF) conformation produces a more rapid rise in accessibility. **F)** 2 TF kinetic model (out of equilibrium) simulations (n=5) measuring binding occupancy of TF A over a range of TF A motif affinities across changing concentrations of TF A. TF B affinity and concentration was kept constant. As concentration of TF A increases, binding occupancy reaches the maximum value of 100%, unlike accessibility state which plateaus at values less than 100% (Figure 5E). Parameter ranges can be found in Table S7. **G)** 2 TF kinetic model (out of equilibrium) simulations (n=5) measuring accessibility cooperativity between TF A and TF B over a range of TF A motif affinities across changing concentrations of TF A. Though TF B affinity and concentration was kept constant, the presence of B caused accessibility states to be cooperatively enhanced. Note that the cooperativity disappears at high TF A concentrations and the maximum cooperativity is increased with affinity. Parameter ranges can be found in Table S7.

**Supplemental Figure 6.**
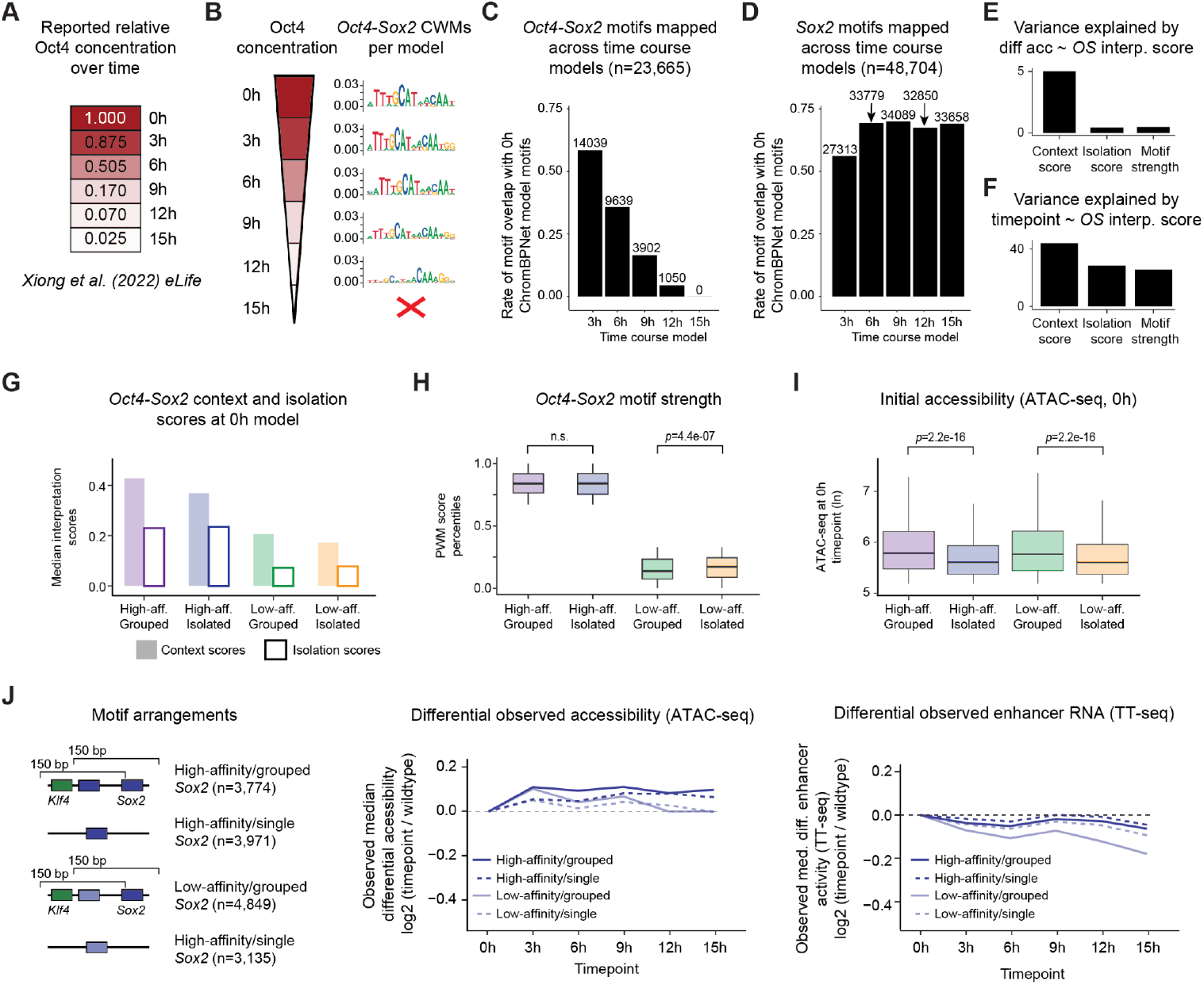
**A)** Reported relative rates of Oct4 upon doxycycline withdrawal time course. **B)** Accessibility TF-MoDISco CWMs of discovered *Oct4-Sox2* motifs from accessibility models of all Oct4 concentration timepoints. Accessibility model capturing the 15h time point failed to discover a contributing role of the *Oct4-Sox2* motif to accessibility. **C-D)** Rates by which **(C)** *Oct4-Sox2* and **(D)** *Sox2* motifs mapped by the reference 0h timepoint model were mapped by the corresponding time course accessibility models. Overall mapping frequencies are reported above each rate. **E)** Barplots measuring the variance explained from interpretation scores to explain *Oct4-Sox2* motif (mapped by the 0h timepoint) effects on observed differential accessibility changes over the concentration time course using independent linear regression. **F)** Barplots measuring the variance explained from interpretation scores to explain *Oct4-Sox2* motif (mapped by the 0h timepoint) effects on which timepoint (treated as a numeric value) each *Oct4-Sox2* motif was last mapped across using independent linear regression for each interpretation method. **G)** Median context and isolation scores of *Oct4-Sox2* motifs mapped by the 0h timepoint model. While both grouped and single motif arrangements possess higher context scores than isolation scores (likely because grouped arrangements only consider *Sox2* and *Klf4* motif partners and not other motifs), grouped arrangements possess a higher relative ratio of the two interpretation scores. **H)** Boxplots depicting PWM score percentiles of *Oct4-Sox2* motifs across the motif arrangements denoted in Figure 6E. Statistics were performed using a Wilcoxson test. **I)** Boxplots depicting initial ATAC-seq reads at the 0h timepoint across the motif arrangements denoted in Figure 6E. Statistics were performed using a Wilcoxson test. Note that grouped motif arrangements possess intrinsically higher initial accessibility levels than single motif arrangements. **J)** Graphic (left) denoting arrangements of single/grouped and strong/weak *Sox2* motifs without an *Oct4-Sox2* motif nearby. Median experimental differential accessibility levels (middle) measured over the concentration time course across regions with either single (solid line) or grouped (dashed line) high- and low-affinity *Sox2* motifs. Median experimental differential enhancer expression levels (right) measured over the concentration time course using TT-seq across regions with either single (solid line) or grouped (dashed line) high- and low-affinity *Sox2* motifs. *Sox2* motifs were derived from the 0h timepoint model, while high- and low-affinity motifs were the top and bottom tertiles of PWM scores.

## References

1. Zeitlinger, J., Roy, S., Ay, F., Mathelier, A., Medina-Rivera, A., Mahony, S., Sinha, S., and Ernst, J. (2025). Perspective on recent developments and challenges in regulatory and systems genomics. Bioinformatics Advances 5, vbaf106. 10.1093/bioadv/vbaf106.

2. Kim, S., and Wysocka, J. (2023). Deciphering the multi-scale, quantitative cis-regulatory code. Mol. Cell 83, 373–392. 10.1016/j.molcel.2022.12.032.

3. Peña-Martínez, E.G., and Rodríguez-Martínez, J.A. (2024). Decoding Non-coding Variants: Recent Approaches to Studying Their Role in Gene Regulation and Human Diseases. Front Biosci (Schol Ed) 16, 4. 10.31083/j.fbs1601004.

4. Farley, E.K., Olson, K.M., Zhang, W., Brandt, A.J., Rokhsar, D.S., and Levine, M.S. (2015). Suboptimization of developmental enhancers. Science 350, 325–328. 10.1126/science.aac6948.

5. Papagianni, A., Forés, M., Shao, W., He, S., Koenecke, N., Andreu, M.J., Samper, N., Paroush, Z., González-Crespo, S., Zeitlinger, J., et al. (2018). Capicua controls Toll/IL-1 signaling targets independently of RTK regulation. Proc Natl Acad Sci USA 115, 1807–1812. 10.1073/pnas.1713930115.

6. Crocker, J., Abe, N., Rinaldi, L., McGregor, A.P., Frankel, N., Wang, S., Alsawadi, A., Valenti, P., Plaza, S., Payre, F., et al. (2015). Low affinity binding site clusters confer hox specificity and regulatory robustness. Cell 160, 191–203. 10.1016/j.cell.2014.11.041.

7. Lim, F., Solvason, J.J., Ryan, G.E., Le, S.H., Jindal, G.A., Steffen, P., Jandu, S.K., and Farley, E.K. (2024). Affinity-optimizing enhancer variants disrupt development. Nature 626, 151–159. 10.1038/s41586-023-06922-8.

8. Peterson, K.A., Nishi, Y., Ma, W., Vedenko, A., Shokri, L., Zhang, X., McFarlane, M., Baizabal, J.-M., Junker, J.P., van Oudenaarden, A., et al. (2012). Neural-specific Sox2 input and differential Gli-binding affinity provide context and positional information in Shh-directed neural patterning. Genes Dev. 26, 2802–2816. 10.1101/gad.207142.112.

9. Jiang, J., and Levine, M. (1993). Binding affinities and cooperative interactions with bHLH activators delimit threshold responses to the dorsal gradient morphogen. Cell 72, 741–752. 10.1016/0092-8674(93)90402-c.

10. Kribelbauer, J.F., Rastogi, C., Bussemaker, H.J., and Mann, R.S. (2019). Low-Affinity Binding Sites and the Transcription Factor Specificity Paradox in Eukaryotes. Annu. Rev. Cell Dev. Biol. 35, 357–379. 10.1146/annurev-cellbio-100617-062719.

11. Wasserman, W.W., and Sandelin, A. (2004). Applied bioinformatics for the identification of regulatory elements. Nat. Rev. Genet. 5, 276–287. 10.1038/nrg1315.

12. Zia, A., and Moses, A.M. (2012). Towards a theoretical understanding of false positives in DNA motif finding. BMC Bioinformatics 13, 151. 10.1186/1471-2105-13-151.

13. Jayaram, N., Usvyat, D., and R Martin, A.C. (2016). Evaluating tools for transcription factor binding site prediction. BMC Bioinformatics 17, 547. 10.1186/s12859-016-1298-9.

14. Stormo, G.D. (2000). DNA binding sites: representation and discovery. Bioinformatics 16, 16–23. 10.1093/bioinformatics/16.1.16.

15. Burz, D.S., Rivera-Pomar, R., Jäckle, H., and Hanes, S.D. (1998). Cooperative DNA-binding by Bicoid provides a mechanism for threshold-dependent gene activation in the Drosophila embryo. EMBO J. 17, 5998–6009. 10.1093/emboj/17.20.5998.

16. Lebrecht, D., Foehr, M., Smith, E., Lopes, F.J.P., Vanario-Alonso, C.E., Reinitz, J., Burz, D.S., and Hanes, S.D. (2005). Bicoid cooperative DNA binding is critical for embryonic patterning in Drosophila. Proc Natl Acad Sci USA 102, 13176–13181. 10.1073/pnas.0506462102.

17. Shahein, A., López-Malo, M., Istomin, I., Olson, E.J., Cheng, S., and Maerkl, S.J. (2022). Systematic analysis of low-affinity transcription factor binding site clusters in vitro and in vivo establishes their functional relevance. Nat. Commun. 13, 5273. 10.1038/s41467-022-32971-0.

18. Barral, A., and Zaret, K.S. (2024). Pioneer factors: roles and their regulation in development. Trends Genet. 40, 134–148. 10.1016/j.tig.2023.10.007.

19. Kim, D.S., Risca, V.I., Reynolds, D.L., Chappell, J., Rubin, A.J., Jung, N., Donohue, L.K.H., Lopez-Pajares, V., Kathiria, A., Shi, M., et al. (2021). The dynamic, combinatorial cis-regulatory lexicon of epidermal differentiation. Nat. Genet. 53, 1564–1576. 10.1038/s41588-021-00947-3.

20. Swinstead, E.E., Miranda, T.B., Paakinaho, V., Baek, S., Goldstein, I., Hawkins, M., Karpova, T.S., Ball, D., Mazza, D., Lavis, L.D., et al. (2016). Steroid Receptors Reprogram FoxA1 Occupancy through Dynamic Chromatin Transitions. Cell 165, 593–605. 10.1016/j.cell.2016.02.067.

21. Brennan, K.J., Weilert, M., Krueger, S., Pampari, A., Liu, H.-Y., Yang, A.W.H., Morrison, J.A., Hughes, T.R., Rushlow, C.A., Kundaje, A., et al. (2023). Chromatin accessibility in the Drosophila embryo is determined by transcription factor pioneering and enhancer activation. Dev. Cell 58, 1898-1916.e9. 10.1016/j.devcel.2023.07.007.

22. Soufi, A., Garcia, M.F., Jaroszewicz, A., Osman, N., Pellegrini, M., and Zaret, K.S. (2015). Pioneer transcription factors target partial DNA motifs on nucleosomes to initiate reprogramming. Cell 161, 555–568. 10.1016/j.cell.2015.03.017.

23. Meers, M.P., Janssens, D.H., and Henikoff, S. (2019). Pioneer Factor-Nucleosome Binding Events during Differentiation Are Motif Encoded. Mol. Cell 75, 562-575.e5. 10.1016/j.molcel.2019.05.025.

24. King, D.M., Hong, C.K.Y., Shepherdson, J.L., Granas, D.M., Maricque, B.B., and Cohen, B.A. (2020). Synthetic and genomic regulatory elements reveal aspects of cis-regulatory grammar in mouse embryonic stem cells. eLife 9. 10.7554/eLife.41279.

25. Hammelman, J., Krismer, K., Banerjee, B., Gifford, D.K., and Sherwood, R.I. (2020). Identification of determinants of differential chromatin accessibility through a massively parallel genome-integrated reporter assay. Genome Res. 30, 1468–1480. 10.1101/gr.263228.120.

26. Pham, T.-H., Minderjahn, J., Schmidl, C., Hoffmeister, H., Schmidhofer, S., Chen, W., Längst, G., Benner, C., and Rehli, M. (2013). Mechanisms of in vivo binding site selection of the hematopoietic master transcription factor PU.1. Nucleic Acids Res. 41, 6391–6402. 10.1093/nar/gkt355.

27. Naqvi, S., Kim, S., Tabatabaee, S., Pampari, A., Kundaje, A., Pritchard, J.K., and Wysocka, J. (2025). Transfer learning reveals sequence determinants of the quantitative response to transcription factor dosage. Cell Genomics 5, 100780. 10.1016/j.xgen.2025.100780.

28. Papatsenko, D., Goltsev, Y., and Levine, M. (2009). Organization of developmental enhancers in the Drosophila embryo. Nucleic Acids Res. 37, 5665–5677. 10.1093/nar/gkp619.

29. Irizarry, J., and Stathopoulos, A. (2021). Dynamic patterning by morphogens illuminated by cis-regulatory studies. Development 148. 10.1242/dev.196113.

30. Driever, W., Thoma, G., and Nüsslein-Volhard, C. (1989). Determination of spatial domains of zygotic gene expression in the Drosophila embryo by the affinity of binding sites for the bicoid morphogen. Nature 340, 363–367. 10.1038/340363a0.

31. Barbadilla-Martínez, L., Klaassen, N., van Steensel, B., and de Ridder, J. (2025). Predicting gene expression from DNA sequence using deep learning models. Nat. Rev. Genet. 10.1038/s41576-025-00841-2.

32. Routhier, E., and Mozziconacci, J. (2022). Genomics enters the deep learning era. PeerJ 10, e13613. 10.7717/peerj.13613.

33. Novakovsky, G., Dexter, N., Libbrecht, M.W., Wasserman, W.W., and Mostafavi, S. (2023). Obtaining genetics insights from deep learning via explainable artificial intelligence. Nat. Rev. Genet. 24, 125–137. 10.1038/s41576-022-00532-2.

34. Alipanahi, B., Delong, A., Weirauch, M.T., and Frey, B.J. (2015). Predicting the sequence specificities of DNA-and RNA-binding proteins by deep learning. Nat. Biotechnol. 33, 831–838. 10.1038/nbt.3300.

35. Dalal, K., McAnany, C., Weilert, M., McKinney, M.C., Krueger, S., and Zeitlinger, J. (2025). Interpreting regulatory mechanisms of Hippo signaling through a deep learning sequence model. Cell Genomics.

36. Avsec, Ž., Weilert, M., Shrikumar, A., Krueger, S., Alexandari, A., Dalal, K., Fropf, R., McAnany, C., Gagneur, J., Kundaje, A., et al. (2021). Base-resolution models of transcription-factor binding reveal soft motif syntax. Nat. Genet. 53, 354–366. 10.1038/s41588-021-00782-6.

37. Liu, B.B., Jessa, S., Kim, S.H., Ng, Y.T., Higashino, S. il, Marinov, G.K., Chen, D.C., Parks, B.E., Li, L., Nguyen, T.C., et al. (2025). Dissecting regulatory syntax in human development with scalable multiomics and deep learning. BioRxiv. 10.1101/2025.04.30.651381.

38. Pampari, A., Shcherbina, A., Kvon, E.Z., Kosicki, M., Nair, S., Kundu, S., Kathiria, A.S., Risca, V.I., Kuningas, K., Alasoo, K., et al. (2025). ChromBPNet: bias factorized, base-resolution deep learning models of chromatin accessibility reveal cis-regulatory sequence syntax, transcription factor footprints and regulatory variants. BioRxiv. 10.1101/2024.12.25.630221.

39. Maslova, A., Ramirez, R.N., Ma, K., Schmutz, H., Wang, C., Fox, C., Ng, B., Benoist, C., Mostafavi, S., and Immunological Genome Project (2020). Deep learning of immune cell differentiation. Proc Natl Acad Sci USA 117, 25655–25666. 10.1073/pnas.2011795117.

40. Agarwal, V., Inoue, F., Schubach, M., Penzar, D., Martin, B.K., Dash, P.M., Keukeleire, P., Zhang, Z., Sohota, A., Zhao, J., et al. (2025). Massively parallel characterization of transcriptional regulatory elements. Nature 639, 411–420. 10.1038/s41586-024-08430-9.

41. de Almeida, B.P., Reiter, F., Pagani, M., and Stark, A. (2022). DeepSTARR predicts enhancer activity from DNA sequence and enables the de novo design of synthetic enhancers. Nat. Genet. 54, 613–624. 10.1038/s41588-022-01048-5.

42. Cochran, K., Yin, M., Mantripragada, A., Schreiber, J., Marinov, G.K., Shah, S.R., Yu, H., Lis, J.T., and Kundaje, A. (2024). Dissecting the cis-regulatory syntax of transcription initiation with deep learning. BioRxiv. 10.1101/2024.05.28.596138.

43. Dudnyk, K., Cai, D., Shi, C., Xu, J., and Zhou, J. (2024). Sequence basis of transcription initiation in the human genome. Science 384, eadj0116. 10.1126/science.adj0116.

44. He, A.Y., and Danko, C.G. (2024). Dissection of core promoter syntax through single nucleotide resolution modeling of transcription initiation. BioRxiv. 10.1101/2024.03.13.583868.

45. Taskiran, I.I., Spanier, K.I., Dickmänken, H., Kempynck, N., Panĉíková, A., Ekşi, E.C., Hulselmans, G., Ismail, J.N., Theunis, K., Vandepoel, R., et al. (2024). Cell-type-directed design of synthetic enhancers. Nature 626, 212–220. 10.1038/s41586-023-06936-2.

46. Yin, C., Castillo-Hair, S., Byeon, G.W., Bromley, P., Meuleman, W., and Seelig, G. (2025). Iterative deep learning design of human enhancers exploits condensed sequence grammar to achieve cell-type specificity. Cell Syst. 16, 101302. 10.1016/j.cels.2025.101302.

47. de Almeida, B.P., Schaub, C., Pagani, M., Secchia, S., Furlong, E.E.M., and Stark, A. (2024). Targeted design of synthetic enhancers for selected tissues in the Drosophila embryo. Nature 626, 207–211. 10.1038/s41586-023-06905-9.

48. Durdu, S., Iskar, M., Isbel, L., Hoerner, L., Wirbelauer, C., Burger, L., Hess, D., Iesmantavicius, V., and Schübeler, D. (2025). Chromatin-dependent motif syntax defines differentiation trajectories. Molecular cell 85, 2900-2918.e16. 10.1016/j.molcel.2025.07.005.

49. Nair, S., Ameen, M., Sundaram, L., Pampari, A., Schreiber, J., Balsubramani, A., Wang, Y.X., Burns, D., Blau, H.M., Karakikes, I., et al. (2023). Transcription factor stoichiometry, motif affinity and syntax regulate single-cell chromatin dynamics during fibroblast reprogramming to pluripotency. BioRxiv. 10.1101/2023.10.04.560808.

50. Arora, S., Yang, J., Akiyama, T., James, D.Q., Morrissey, A., Blanda, T.R., Badjatia, N., Lai, W.K.M., Ko, M.S.H., Pugh, B.F., et al. (2023). Joint sequence & chromatin neural networks characterize the differential abilities of Forkhead transcription factors to engage inaccessible chromatin. BioRxiv. 10.1101/2023.10.06.561228.

51. McAnany, C., Weilert, M., Kamulegeya, F., Mehta, G., Gardner, J.M., Kundaje, A., and Zeitlinger, J. (2025). PISA: a versatile interpretation tool for visualizing cis-regulatory rules in genomic data. BioRxiv.

52. Shrikumar, A., Greenside, P., and Kundaje, A. (2017). Learning Important Features Through Propagating Activation Differences. In (Proceedings of Machine Learning Research), pp. 3145–3153.

53. Lundberg, S.M., and Lee, S.-I. (2017). A Unified Approach to Interpreting Model Predictions. Advances in Neural Information Processing Systems.

54. Shrikumar, A., Tian, K., Avsec, Z., Shcherbina, A., Banerjee, A., Sharmin, M., Nair, S., and Kundaje, A. (2020). Technical Note on Transcription Factor Motif Discovery from Importance Scores (TF-MoDISco) version 0.5.6.5. arXiv.

55. Soufi, A., Donahue, G., and Zaret, K.S. (2012). Facilitators and impediments of the pluripotency reprogramming factors’ initial engagement with the genome. Cell 151, 994–1004. 10.1016/j.cell.2012.09.045.

56. Di Giammartino, D.C., Kloetgen, A., Polyzos, A., Liu, Y., Kim, D., Murphy, D., Abuhashem, A., Cavaliere, P., Aronson, B., Shah, V., et al. (2019). KLF4 is involved in the organization and regulation of pluripotency-associated three-dimensional enhancer networks. Nat. Cell Biol. 21, 1179–1190. 10.1038/s41556-019-0390-6.

57. Buecker, C., Srinivasan, R., Wu, Z., Calo, E., Acampora, D., Faial, T., Simeone, A., Tan, M., Swigut, T., and Wysocka, J. (2014). Reorganization of enhancer patterns in transition from naive to primed pluripotency. Cell Stem Cell 14, 838–853. 10.1016/j.stem.2014.04.003.

58. Maresca, M., van den Brand, T., Li, H., Teunissen, H., Davies, J., and de Wit, E. (2023). Pioneer activity distinguishes activating from non-activating SOX2 binding sites. EMBO J. 42, e113150. 10.15252/embj.2022113150.

59. Grant, C.E., Bailey, T.L., and Noble, W.S. (2011). FIMO: scanning for occurrences of a given motif. Bioinformatics 27, 1017–1018. 10.1093/bioinformatics/btr064.

60. Bailey, T.L., Boden, M., Buske, F.A., Frith, M., Grant, C.E., Clementi, L., Ren, J., Li, W.W., and Noble, W.S. (2009). MEME SUITE: tools for motif discovery and searching. Nucleic Acids Res. 37, W202–8. 10.1093/nar/gkp335.

61. Lai, W.K.M., Mariani, L., Rothschild, G., Smith, E.R., Venters, B.J., Blanda, T.R., Kuntala, P.K., Bocklund, K., Mairose, J., Dweikat, S.N., et al. (2021). A ChIP-exo screen of 887 Protein Capture Reagents Program transcription factor antibodies in human cells. Genome Res. 31, 1663–1679. 10.1101/gr.275472.121.

62. Mariani, L., Weinand, K., Vedenko, A., Barrera, L.A., and Bulyk, M.L. (2017). Identification of Human Lineage-Specific Transcriptional Coregulators Enabled by a Glossary of Binding Modules and Tunable Genomic Backgrounds. Cell Syst. 5, 187-201.e7. 10.1016/j.cels.2017.06.015.

63. Badis, G., Berger, M.F., Philippakis, A.A., Talukder, S., Gehrke, A.R., Jaeger, S.A., Chan, E.T., Metzler, G., Vedenko, A., Chen, X., et al. (2009). Diversity and complexity in DNA recognition by transcription factors. Science 324, 1720–1723. 10.1126/science.1162327.

64. Newburger, D.E., and Bulyk, M.L. (2009). UniPROBE: an online database of protein binding microarray data on protein-DNA interactions. Nucleic Acids Res. 37, D77–82. 10.1093/nar/gkn660.

65. Waite, J.B., Boytz, R., Traeger, A.R., Lind, T.M., Lumbao-Conradson, K., and Torigoe, S.E. (2024). A suboptimal OCT4-SOX2 binding site facilitates the naïve-state specific function of a Klf4 enhancer. PLoS ONE 19, e0311120. 10.1371/journal.pone.0311120.

66. Srivastava, D., and Mahony, S. (2020). Sequence and chromatin determinants of transcription factor binding and the establishment of cell type-specific binding patterns. Biochim. Biophys. Acta Gene Regul. Mech. 1863, 194443. 10.1016/j.bbagrm.2019.194443.

67. Xiong, L., Tolen, E.A., Choi, J., Velychko, S., Caizzi, L., Velychko, T., Adachi, K., MacCarthy, C.M., Lidschreiber, M., Cramer, P., et al. (2022). Oct4 differentially regulates chromatin opening and enhancer transcription in pluripotent stem cells. eLife 11. 10.7554/eLife.71533.

68. Alexandari, A.M., Horton, C.A., Shrikumar, A., Shah, N., Li, E., Weilert, M., Pufall, M.A., Zeitlinger, J., Fordyce, P.M., and Kundaje, A. (2023). De novo distillation of thermodynamic affinity from deep learning regulatory sequence models of in vivo protein-DNA binding. BioRxiv. 10.1101/2023.05.11.540401.

69. Koo, P.K., Majdandzic, A., Ploenzke, M., Anand, P., and Paul, S.B. (2021). Global importance analysis: An interpretability method to quantify importance of genomic features in deep neural networks. PLoS Comput. Biol. 17, e1008925. 10.1371/journal.pcbi.1008925.

70. Kelley, D.R., Snoek, J., and Rinn, J.L. (2016). Basset: learning the regulatory code of the accessible genome with deep convolutional neural networks. Genome Res. 26, 990–999. 10.1101/gr.200535.115.

71. Nair, S., Shrikumar, A., Schreiber, J., and Kundaje, A. (2022). fastISM: performant in silico saturation mutagenesis for convolutional neural networks. Bioinformatics 38, 2397–2403. 10.1093/bioinformatics/btac135.

72. Toneyan, S., and Koo, P.K. (2024). Interpreting cis-regulatory interactions from large-scale deep neural networks. Nat. Genet. 56, 2517–2527. 10.1038/s41588-024-01923-3.

73. Xie, L., Torigoe, S.E., Xiao, J., Mai, D.H., Li, L., Davis, F.P., Dong, P., Marie-Nelly, H., Grimm, J., Lavis, L., et al. (2017). A dynamic interplay of enhancer elements regulates Klf4 expression in naïve pluripotency. Genes Dev. 31, 1795–1808. 10.1101/gad.303321.117.

74. King, H.W., and Klose, R.J. (2017). The pioneer factor OCT4 requires the chromatin remodeller BRG1 to support gene regulatory element function in mouse embryonic stem cells. eLife 6. 10.7554/eLife.22631.

75. Gibson, T.J., Larson, E.D., and Harrison, M.M. (2024). Protein-intrinsic properties and context-dependent effects regulate pioneer factor binding and function. Nat. Struct. Mol. Biol. 31, 548–558. 10.1038/s41594-024-01231-8.

76. Martin, V., Zhuang, F., Zhang, Y., Pinheiro, K., and Gordân, R. (2023). High-throughput data and modeling reveal insights into the mechanisms of cooperative DNA-binding by transcription factor proteins. Nucleic Acids Res. 51, 11600–11612. 10.1093/nar/gkad872.

77. Mirny, L.A. (2010). Nucleosome-mediated cooperativity between transcription factors. Proc Natl Acad Sci USA 107, 22534–22539. 10.1073/pnas.0913805107.

78. Sönmezer, C., Kleinendorst, R., Imanci, D., Barzaghi, G., Villacorta, L., Schübeler, D., Benes, V., Molina, N., and Krebs, A.R. (2021). Molecular Co-occupancy Identifies Transcription Factor Binding Cooperativity In Vivo. Mol. Cell 81, 255-267.e6. 10.1016/j.molcel.2020.11.015.

79. Doughty, B.R., Hinks, M.M., Schaepe, J.M., Marinov, G.K., Thurm, A.R., Rios-Martinez, C., Parks, B.E., Tan, Y., Marklund, E., Dubocanin, D., et al. (2024). Single-molecule states link transcription factor binding to gene expression. Nature 636, 745–754. 10.1038/s41586-024-08219-w.

80. Adams, C.C., and Workman, J.L. (1995). Binding of disparate transcriptional activators to nucleosomal DNA is inherently cooperative. Mol. Cell. Biol. 15, 1405–1421. 10.1128/MCB.15.3.1405.

81. He, X., Samee, M.A.H., Blatti, C., and Sinha, S. (2010). Thermodynamics-based models of transcriptional regulation by enhancers: the roles of synergistic activation, cooperative binding and short-range repression. PLoS Comput. Biol. 6. 10.1371/journal.pcbi.1000935.

82. Giorgetti, L., Siggers, T., Tiana, G., Caprara, G., Notarbartolo, S., Corona, T., Pasparakis, M., Milani, P., Bulyk, M.L., and Natoli, G. (2010). Noncooperative interactions between transcription factors and clustered DNA binding sites enable graded transcriptional responses to environmental inputs. Mol. Cell 37, 418–428. 10.1016/j.molcel.2010.01.016.

83. Kharerin, H., Bhat, P.J., Marko, J.F., and Padinhateeri, R. (2016). Role of transcription factor-mediated nucleosome disassembly in PHO5 gene expression. Sci. Rep. 6, 20319. 10.1038/srep20319.

84. Wolff, M.R., Schmid, A., Korber, P., and Gerland, U. (2021). Effective dynamics of nucleosome configurations at the yeast PHO5 promoter. eLife 10. 10.7554/eLife.58394.

85. Strebinger, D., Deluz, C., Friman, E.T., Govindan, S., Alber, A.B., and Suter, D.M. (2019). Endogenous fluctuations of OCT4 and SOX2 bias pluripotent cell fate decisions. Mol. Syst. Biol. 15, e9002. 10.15252/msb.20199002.

86. Malaga Gadea, F.C., and Nikolova, E.N. (2023). Structural Plasticity of Pioneer Factor Sox2 and DNA Bendability Modulate Nucleosome Engagement and Sox2-Oct4 Synergism. J. Mol. Biol. 435, 167916. 10.1016/j.jmb.2022.167916.

87. Chen, J., Zhang, Z., Li, L., Chen, B.-C., Revyakin, A., Hajj, B., Legant, W., Dahan, M., Lionnet, T., Betzig, E., et al. (2014). Single-molecule dynamics of enhanceosome assembly in embryonic stem cells. Cell 156, 1274–1285. 10.1016/j.cell.2014.01.062.

88. Hemphill, W.O., Steiner, H.R., Kominsky, J.R., Wuttke, D.S., and Cech, T.R. (2024). Transcription factors ERα and Sox2 have differing multiphasic DNA- and RNA-binding mechanisms. RNA 30, 1089–1105. 10.1261/rna.080027.124.

89. Marinov, G.K., Doughty, B.R., Schaepe, J.M., Wang, T., Smith, M.M., Chen, M., Kundaje, A., Sun, Z., and Greenleaf, W. (2025). The human transcription factor occupancy landscape viewed using high-resolution in situ base-conversion strand-specific single-molecule chromatin accessibility mapping. BioRxiv. 10.1101/2025.06.27.662080.

90. Love, M.I., Huber, W., and Anders, S. (2014). Moderated estimation of fold change and dispersion for RNA-seq data with DESeq2. Genome Biol. 15, 550. 10.1186/s13059-014-0550-8.

91. Villar, D., Flicek, P., and Odom, D.T. (2014). Evolution of transcription factor binding in metazoans - mechanisms and functional implications. Nat. Rev. Genet. 15, 221–233. 10.1038/nrg3481.

92. Fuqua, T., Jordan, J., van Breugel, M.E., Halavatyi, A., Tischer, C., Polidoro, P., Abe, N., Tsai, A., Mann, R.S., Stern, D.L., et al. (2020). Dense encoding of developmental regulatory information may constrain evolvability. BioRxiv. 10.1101/2020.04.17.046052.

93. Vincent, B.J., Estrada, J., and DePace, A.H. (2016). The appeasement of Doug: a synthetic approach to enhancer biology. Integr Biol (Camb) 8, 475–484. 10.1039/c5ib00321k.

94. Ling, L., Mühling, B., Jaenichen, R., and Gompel, N. (2023). Increased chromatin accessibility promotes the evolution of a transcriptional silencer in Drosophila. Sci. Adv. 9, eade6529. 10.1126/sciadv.ade6529.

95. Montavon, T., Soshnikova, N., Mascrez, B., Joye, E., Thevenet, L., Splinter, E., de Laat, W., Spitz, F., and Duboule, D. (2011). A regulatory archipelago controls Hox genes transcription in digits. Cell 147, 1132–1145. 10.1016/j.cell.2011.10.023.

96. Yamada, S., Whitney, P.H., Huang, S.-K., Eck, E.C., Garcia, H.G., and Rushlow, C.A. (2019). The drosophila pioneer factor zelda modulates the nuclear microenvironment of a dorsal target enhancer to potentiate transcriptional output. Curr. Biol. 29, 1387-1393.e5. 10.1016/j.cub.2019.03.019.

97. Foo, S.M., Sun, Y., Lim, B., Ziukaite, R., O’Brien, K., Nien, C.-Y., Kirov, N., Shvartsman, S.Y., and Rushlow, C.A. (2014). Zelda potentiates morphogen activity by increasing chromatin accessibility. Curr. Biol. 24, 1341–1346. 10.1016/j.cub.2014.04.032.

98. Keller, S.H., Jena, S.G., Yamazaki, Y., and Lim, B. (2020). Regulation of spatiotemporal limits of developmental gene expression via enhancer grammar. Proc Natl Acad Sci USA 117, 15096–15103. 10.1073/pnas.1917040117.

99. Zhu, F., Farnung, L., Kaasinen, E., Sahu, B., Yin, Y., Wei, B., Dodonova, S.O., Nitta, K.R., Morgunova, E., Taipale, M., et al. (2018). The interaction landscape between transcription factors and the nucleosome. Nature 562, 76–81. 10.1038/s41586-018-0549-5.

100. Makowski, M.M., Gaullier, G., and Luger, K. (2020). Picking a nucleosome lock: Sequence- and structure-specific recognition of the nucleosome. J. Biosci. 45. 10.1007/s12038-019-9970-7.

101. Michael, A.K., Grand, R.S., Isbel, L., Cavadini, S., Kozicka, Z., Kempf, G., Bunker, R.D., Schenk, A.D., Graff-Meyer, A., Pathare, G.R., et al. (2020). Mechanisms of OCT4-SOX2 motif readout on nucleosomes. Science.

102. Boeger, H., Griesenbeck, J., and Kornberg, R.D. (2008). Nucleosome retention and the stochastic nature of promoter chromatin remodeling for transcription. Cell 133, 716–726. 10.1016/j.cell.2008.02.051.

103. Donovan, B.T., Luo, Y., Meng, Z., and Poirier, M.G. (2023). The nucleosome unwrapping free energy landscape defines distinct regions of transcription factor accessibility and kinetics. Nucleic Acids Res. 51, 1139–1153. 10.1093/nar/gkac1267.

104. Scholes, C., DePace, A.H., and Sánchez, Á. (2017). Combinatorial Gene Regulation through Kinetic Control of the Transcription Cycle. Cell Syst. 4, 97-108.e9. 10.1016/j.cels.2016.11.012.

105. Avsec, Ž., Agarwal, V., Visentin, D., Ledsam, J.R., Grabska-Barwinska, A., Taylor, K.R., Assael, Y., Jumper, J., Kohli, P., and Kelley, D.R. (2021). Effective gene expression prediction from sequence by integrating long-range interactions. Nat. Methods 18, 1196–1203. 10.1038/s41592-021-01252-x.

106. Karollus, A., Mauermeier, T., and Gagneur, J. (2023). Current sequence-based models capture gene expression determinants in promoters but mostly ignore distal enhancers. Genome Biol. 24, 56. 10.1186/s13059-023-02899-9.

107. Huang, C., Shuai, R.W., Baokar, P., Chung, R., Rastogi, R., Kathail, P., and Ioannidis, N.M. (2023). Personal transcriptome variation is poorly explained by current genomic deep learning models. Nat. Genet. 55, 2056–2059. 10.1038/s41588-023-01574-w.

108. Connelly, J.P., and Pruett-Miller, S.M. (2019). CRIS.py: A Versatile and High-throughput Analysis Program for CRISPR-based Genome Editing. Sci. Rep. 9, 4194. 10.1038/s41598-019-40896-w.

109. Corces, M.R., Trevino, A.E., Hamilton, E.G., Greenside, P.G., Sinnott-Armstrong, N.A., Vesuna, S., Satpathy, A.T., Rubin, A.J., Montine, K.S., Wu, B., et al. (2017). An improved ATAC-seq protocol reduces background and enables interrogation of frozen tissues. Nat. Methods 14, 959–962. 10.1038/nmeth.4396.

110. He, Q., Johnston, J., and Zeitlinger, J. (2015). ChIP-nexus enables improved detection of in vivo transcription factor binding footprints. Nat. Biotechnol. 33, 395–401. 10.1038/nbt.3121.

111. Martin, M. (2011). Cutadapt removes adapter sequences from high-throughput sequencing reads. EMBnet j. 17, 10. 10.14806/ej.17.1.200.

112. Langmead, B., and Salzberg, S.L. (2012). Fast gapped-read alignment with Bowtie 2. Nat. Methods 9, 357–359. 10.1038/nmeth.1923.

113. Broad Institute Picard Tools. http://broadinstitute.github.io/picard.

114. Zhang, Y., Liu, T., Meyer, C.A., Eeckhoute, J., Johnson, D.S., Bernstein, B.E., Nusbaum, C., Myers, R.M., Brown, M., Li, W., et al. (2008). Model-based analysis of ChIP-Seq (MACS). Genome Biol. 9, R137. 10.1186/gb-2008-9-9-r137.

115. Li, Q., Brown, J.B., Huang, H., and Bickel, P.J. (2011). Measuring reproducibility of high-throughput experiments. Ann. Appl. Stat. 5, 1752–1779. 10.1214/11-AOAS466.

116. Cleveland, W.S., and Devlin, S.J. (1988). Locally weighted regression: an approach to regression analysis by local fitting. J. Am. Stat. Assoc. 83, 596–610. 10.1080/01621459.1988.10478639.

117. Wickham, H. (2016). ggplot2: Elegant Graphics for Data Analysis | SpringerLink. https://link.springer.com/book/10.1007/978-3-319-24277-4?trk=public_post_comment-text.

118. Dobin, A., Davis, C.A., Schlesinger, F., Drenkow, J., Zaleski, C., Jha, S., Batut, P., Chaisson, M., and Gingeras, T.R. (2013). STAR: ultrafast universal RNA-seq aligner. Bioinformatics 29, 15–21. 10.1093/bioinformatics/bts635.

119. Harrison, P.W., Amode, M.R., Austine-Orimoloye, O., Azov, A.G., Barba, M., Barnes, I., Becker, A., Bennett, R., Berry, A., Bhai, J., et al. (2024). Ensembl 2024. Nucleic Acids Res. 52, D891–D899. 10.1093/nar/gkad1049.

120. Saxonov, S., Berg, P., and Brutlag, D.L. (2006). A genome-wide analysis of CpG dinucleotides in the human genome distinguishes two distinct classes of promoters. Proc Natl Acad Sci USA 103, 1412–1417. 10.1073/pnas.0510310103.

121. Estrada, J., Wong, F., DePace, A., and Gunawardena, J. (2016). Information integration and energy expenditure in gene regulation. Cell 166, 234–244. 10.1016/j.cell.2016.06.012.

122. Gunawardena, J. (2012). A linear framework for time-scale separation in nonlinear biochemical systems. PLoS ONE 7, e36321. 10.1371/journal.pone.0036321.

123. Virtanen, P., Gommers, R., Oliphant, T.E., Haberland, M., Reddy, T., Cournapeau, D., Burovski, E., Peterson, P., Weckesser, W., Bright, J., et al. (2020). SciPy 1.0: fundamental algorithms for scientific computing in Python. Nat. Methods 17, 261–272. 10.1038/s41592-019-0686-2.

124. Mahdavi, S.D., Salmon, G.L., Daghlian, P., Garcia, H.G., and Phillips, R. (2024). Flexibility and sensitivity in gene regulation out of equilibrium. Proc Natl Acad Sci USA 121, e2411395121. 10.1073/pnas.2411395121.

